# Assessment of tobacco and *N. benthamiana* as biofactories of irregular monoterpenes for sustainable crop protection

**DOI:** 10.1101/2023.08.02.551635

**Authors:** Rubén Mateos-Fernández, Sandra Vacas, Ismael Navarro-Fuertes, Vicente Navarro-Llopis, Diego Orzáez, Silvia Gianoglio

## Abstract

Irregular monoterpenes are important precursors of different compounds employed in pest control such as insecticides and insect sex pheromones. Metabolically engineered plants are appealing as biofactories of such compounds, but specially as potential live biodispensers of related bioactive volatiles, which could be continuously emitted to the environment from different plant tissues. Here we assess the use of cultivated tobacco and Nicotiana benthamiana as biofactories for the irregular monoterpenes chrysanthemol and lavandulol. We evaluate the impact of high levels of constitutive metabolite production on the plant physiology and biomass, and their biosynthetic dynamics for different plant tissues and developmental stages. As an example of an active pheromone compound, we super-transformed the best lavandulol-producing tobacco line with an acetyl transferase gene to obtain a tobacco lavandulyl acetate biodispenser emitting up to 0.63 mg of lavandulyl acetate per plant every day. We estimate that with these volatile emission levels, between 200 and 500 plants per hectare would be sufficient to ensure a daily emission of pheromones comparable to commercial lures. This is an important step towards plant-based sustainable solutions for pest control, and it lays the ground for further developing biofactories for other irregular monoterpenoid pheromones, whose biosynthetic genes are yet unknown.

## Introduction

Sustainability in agri-food systems, as in any other sector of the economy, is achieved by balancing the long-term – and sometimes competing – interests of environmental protection, economic profitability and social equity. The protection of crops and stored goods from damage induced by insect pests is an indispensable aspect of increasing agricultural yields and reducing food waste, and the way in which pest control is achieved is fundamental for sustainability. Insect pheromones are sustainable alternatives to traditional pesticides because they are effective, safe, and pose very limited risks of insurgence of genetic resistance, with virtually inexistent effects on non-target populations (Rizvi *et al*., 2021). Pheromones are volatile organic compounds (VOCs) produced by insects that function at very low concentrations as semiochemicals modulating the behavior of conspecifics. The most interesting for pest control are sex pheromones, usually produced by females, and aggregation pheromones, mainly produced by males to attract both males and females. Both types of pheromones can be used as lures to construct traps to attract and affect individuals, or to monitor population levels. Sex pheromones are also employed in mating disruption strategies, where the release of the pheromone to the environment masks the signal produced by females, preventing or delaying mating (Miller & Gut, 2015). Despite their advantages, the chemical synthesis of pheromone compounds and the formulation of traps can be complex and costly, making them affordable for expensive end products (like high-value orchard productions), but far less accessible for row crops (Bento *et al*., 2016; Ioriatti & Lucchi, 2016; Petkevicius *et al*., 2020). Thus, research has focused on developing pheromone biofactories through the engineering of biological hosts like yeasts and plants (Mateos Fernández *et al*., 2022). Bioproduction of pheromones has, in principle, several advantages over chemical synthesis: renewable feedstocks benefit the production pipeline and generate fewer polluting by-products; production costs are reduced; finally, chemical synthesis produces racemic mixtures, while enzymes ensure stereoselectivity, which is crucial for pheromone activity (Mateos Fernández *et al*., 2022). Biofactories can be used either to synthesize active pheromone compounds (Ding *et al*., 2014; Holkenbrink *et al*., 2020; Mateos-Fernández *et al*., 2021), or to produce precursors to be extracted and modified chemically, resulting in hemisynthetic preparations that still enhance sustainability (Nešněrová *et al*., 2004; Xia *et al*., 2020; Wang *et al*., 2022). While yeasts can provide greater yields and ease of extraction, plants may be used as pheromone biofactories following two different strategies: one is to synthesize molecules (active compounds or precursors) to be extracted; the second is to engineer plants to be live pheromone emitters (Bruce *et al*., 2015). The appropriate plant host for each strategy depends on plant biomass, on specialized metabolisms supporting the production of target compounds and, for bioemitters, on the ability to volatilize them.

The bulk of pheromone bioproduction has focused on Lepidopteran sex pheromones, because of the enormous economic relevance of these pests and because these molecules have relatively simple structures and their biosynthetic pathways are known (Löfstedt *et al*., 2016). Still, an immense potential exists to produce a wide variety of pheromones for different targets. Mealybugs (Pseudococcidae) are a family of insects which constitute a relevant threat to crops in sub-tropical and Mediterranean climates. Their mating behavior strongly depends on sex pheromones: these typically contain various monoterpene-derived esters and many species synthesize irregular monoterpenes, which are unusual in nature, resulting from the non-head-to-tail coupling of two DMAPP units instead of the regular (head-to-tail) condensation of an IPP and a DMAPP unit (Kobayashi & Kuzuyama, 2019). Zou & Millar (2015) provide an extensive review of mealybug sex pheromones and their chemistry. Unfortunately, their biosynthesis remains unclear and insect genes responsible for their production are yet to be identified (Tabata, 2022; Juteršek *et al*., 2023). In the absence of known mealybug biosynthetic genes, alternative approaches to the bioproduction of their sex pheromones rely on other organisms producing analogous compounds. This is the case of plants producing lavandulyl pyrophosphate (LPP) and chrysanthemyl pyrophosphate (CPP), irregular branched and cyclic monoterpenoids, respectively (Figure 1A). LPP and its derivatives lavandulol and lavandulyl acetate are produced by various lavender species (Lamiaceae) and by some Apiaceae and are important fragrances, while CPP and the alcohol chrysanthemol are produced by members of the Anthemidae tribe within the Asteraceae family (Minteguiaga *et al*., 2023). LPP and CPP are the precursors of a variety of bioactive compounds important for pest management. Both LPP and CPP are valuable as the monoterpene moieties of the sex pheromone compounds of various mealybug species (such as *Planococcus ficus* Signoret, *Dysmicoccus grassii* Leonardi and *Phenacoccus madeirensis* Green, among others) and can be easily esterified to give an active product (Zou & Millar, 2015). In particular, lavandulyl acetate is an active pheromone compound for the mealybug *D. grassii* (De Alfonso *et al*., 2012), and a component of the aggregation pheromone of the Western flower thrips *Frankliniella occidentalis* Pergrande (Hamilton *et al*., 2005). Finally, lavandulyl acetate has also been identified as a mosquito larvicide with low toxicity towards non-target organisms (Govindarajan & Benelli, 2016), and CPP is a precursor for the biosynthesis of pyrethrins, a class of important natural insecticides (Xu *et al*., 2018).

**Figure 1.**
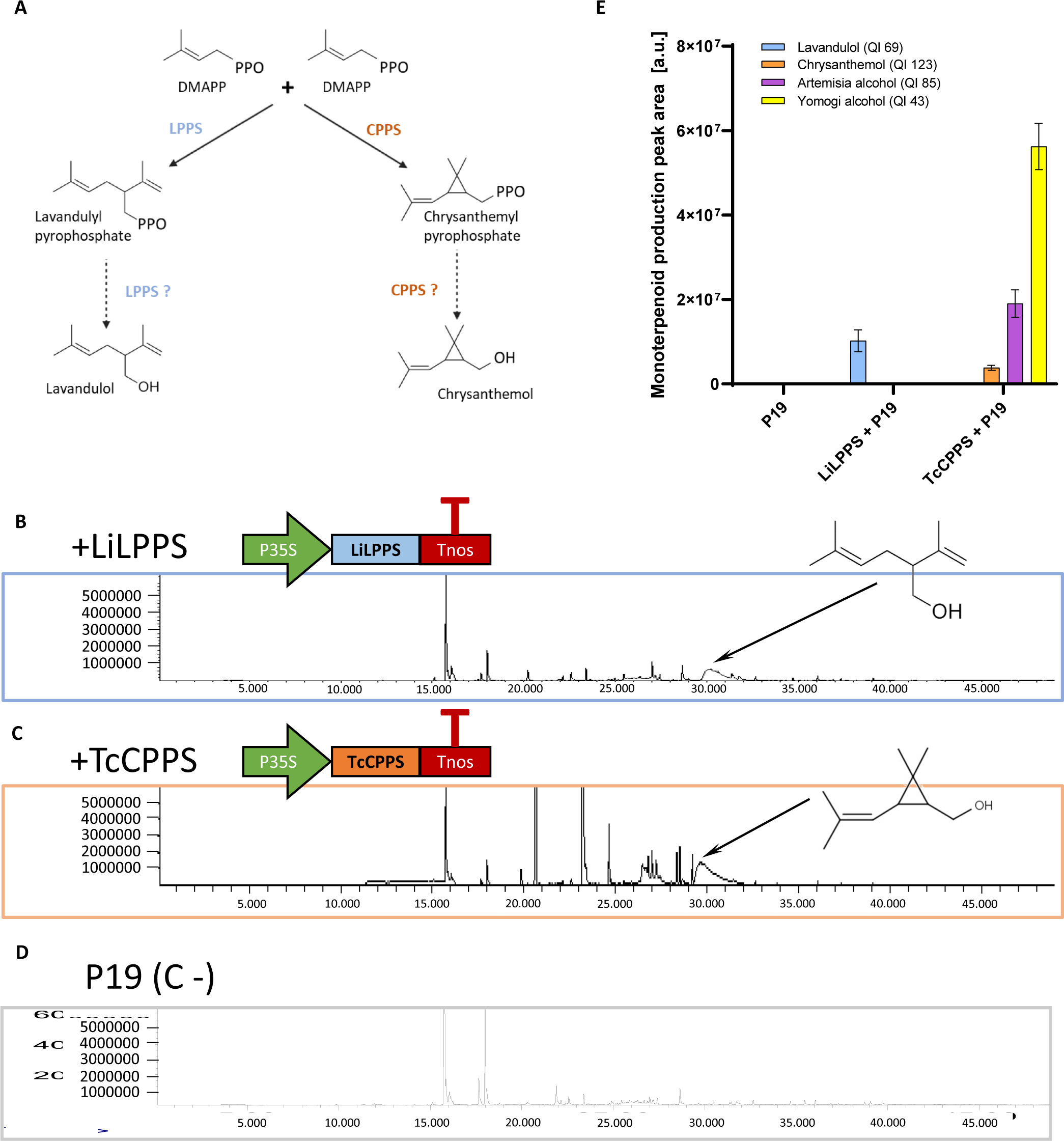
Production of the volatile monoterpenoids lavandulol and chrysanthemol in transient expression in *N. benthamiana*. **(A)** Biosynthetic metabolic pathway of the irregular monoterpenoids lavandulyl pyrophosphate and chrysanthemyl pyrophosphate via the non-head-to-tail condensation of two DMAPP by the LPPS or CPPS enzyme. Production of the alcohols might be due to host endogenous phosphatases or to a bifunctional activity of these irregular IDSs. **(B)** The T-DNA construct used for transient expression of *Li*LPPS controlled by the CaMV35S promoter and Nos terminator, and the GC-MS profile of *N. benthamiana* leaf tissue 5 dpi. **(C)** The T-DNA construct used for transient expression of *Tc*CPPS controlled by the CaMV35S promoter and Nos terminator, and the GC-MS profile of *N. benthamiana* leaf tissue 5 dpi. **(D)** The GC-MS profile of *N. benthamiana* leaf tissue 5 dpi in negative control plants infiltrated only with the P19 silencing suppressor. **(E)** Lavandulol, chrysanthemol, and chrysanthemol-derived compounds measured by GC-MS in agroinfiltrated *N. benthamiana* leaves.

Monoterpenoids are highly accumulated by common aromatic plants, which store essential oil compounds in glandular trichomes. However, these plants are not ideal bioproduction platforms for heterologous compounds, since they are not easy to transform genetically, and their biomass and growth rate are lower than other wide-leaf species, such as tobacco (*Nicotiana tabacum* L.) and *Nicotiana benthamiana* Domin. Tobacco represents a versatile chassis for genetic manipulation with high biomass production, and *N. benthamiana* allows efficient testing of multiple gene combinations through agroinfiltration (Molina-Hidalgo *et al*., 2021). Thus, we used the LPP synthase gene from *Lavandula* × *intermedia* Emeric ex Loisel. (*LiLPPS*; Demissie *et al*., 2013) and the CPP synthase gene from *Tanacetum cinerariifolium* (Trevir.) Sch.Bip. (*TcCPPS*; Yang *et al*., 2014) to transform tobacco and *N. benthamiana*, aiming at assessing the potential of these species as producers and emitters of irregular monoterpenoids. We also transformed *Li*LPPS-expressing tobacco plants with the AAT4 acetyltransferase from *L. intermedia* (Sarker & Mahmoud, 2015), successfully esterifying lavandulol to lavandulyl acetate.

## Materials and methods

### DNA assembly and cloning

All DNA parts used for plant transformation were domesticated and assembled using the GoldenBraid standard as described by Sarrion-Perdigones *et al*. (2011). All constructs were verified by Sanger sequencing and/or restriction analysis. All GB constructs designed and employed in this study are available at www.gbcloning.upv.es under their corresponding IDs, which are listed in Supplementary Table S1. All constructs were cloned using the *Escherichia coli* TOP 10 strain. The final expression vectors were transformed into electrocompetent *Agrobacterium tumefaciens* GV3101 or LBA4404 for transient or stable transformations, respectively.

### Transient expression assays in *Nicotiana benthamiana*

Transient expression assays to validate gene activity were carried out through infiltration of *N. benthamiana* leaves mediated by *Agrobacterium tumefaciens*. Pre-cultures were grown from glycerol stocks for two days at 28°C at 250 rpm with the appropriate antibiotics until saturation, then refreshed and grown overnight in the same conditions. Cells were pelleted and resuspended in an agroinfiltration buffer containing 10 mM 2-(N-morpholino) ethanesulfonic acid (MES), pH 5.7, 10 mM MgCl_2_, and 200 µM acetosyringone, then incubated for 2h at RT under slow shaking. The OD_600_ of each culture was adjusted to reach a value of 0.05-0.06 in the final culture mixtures. Each mixture had a final OD_600_ value of 0.2. Equal volumes of each culture were mixed, including the silencing suppressor P19 for co-infiltration to reduce post-transcriptional gene silencing (Garabagi *et al*., 2012). The relative abundance of each *A. tumefaciens* culture was kept constant in all infiltration mixtures by adding an *A. tumefaciens* culture carrying an empty vector when needed. Agroinfiltration was carried out with a 1 mL needle-free syringe, through the abaxial surface of the three youngest fully expanded leaves of 4-5 weeks old *N. benthamiana* plants, grown at 24°C (light)/20°C (darkness) with a 16:8 h light:darkness photoperiod. Samples were collected 5 days post-agroinfiltration using a Ø 1.5-2 cm corkborer and snap frozen in liquid nitrogen.

### Generation and selection of stable transformants

Stable transgenic plants were generated following the transformation protocol described by Kallam *et al*. (2023). The same procedure was used for tobacco and *N. benthamiana*. For the selection of transgenic progenies, seeds were disinfected by incubation under suspension in 10% trisodium phosphate dodecahydrate for 20’ and then in 3% sodium hypochlorite for 20’. Then, seeds were washed in sterile distilled water and sown on germination medium (5 g/L MS with vitamins, 30 g/L sucrose, 9 g/L Phytoagar, pH = 5.7) supplemented with 100 mg/L kanamycin for positive transgene selection. *NtLPPS-AAT4* tobacco seeds were supplemented with both 100 mg/L kanamycin and 20 mg/L hygromycin for simultaneous transgenic selection. Control plants were obtained similarly, by placing seeds on a non-selective germination medium. WT and antibiotic-resistant seedlings were transferred to the greenhouse 15 days after germination, where they were grown at 24 : 20°C (light : darkness) with a 16 : 8 h light :darkness photoperiod. The segregation of transgenes and the estimation of transgene copy number were determined by calculating survival percentages of seedlings germinated on selective media. Transgene copy number was estimated using the Chi squared test. T_0_ lines were assumed to be multiple copy lines for the transgene when segregation of the transgene was not possible in the T_1_ progeny, and no segregation was detected in the following transgenic generations.

### Plant sampling

Samples for VOCs analysis in the T_0_, T_1_ and T_2_ generations were collected from the youngest fully expanded leaves of 35-40 days-old *N. benthamiana* or *N. tabacum* plants using a Ø 1.5-2 cm corkborer and snap frozen in liquid nitrogen.

For the analysis of T_3_ *N. benthamiana* plants, the first collection of leaf tissue was performed just before the first flower reached anthesis (-1 day), choosing the youngest leaves ranging in length from 3 to 5 cm (henceforth named young leaves), and middle-stem fully expanded leaves (henceforth named adult leaves). The second collection of leaf tissue (post-flowering stage) was carried out after 90 days in soil, following the same criteria adopted in the first collection for the selection of young and adult leaves. For senescent leaves, leaves in the lower part of the stem were sampled when turning slightly yellow. Three leaves per leaf type were sampled as biological replicates.

For the analysis of T_3_ *NtLPPS* and *NtCPPS* tobacco plants, and of T_1_ *NtLPPS-AAT4* tobacco plants, leaf tissue collection was performed just before the first flower reached anthesis (-1 day), sampling upper-stem leaves ranging in length from 15 to 25 cm (henceforth named young leaves), middle-stem fully expanded and deep green leaves (henceforth named adult leaves), and lower-stem leaves, turning slightly yellow, for senescent leaves. Three leaves per leaf type were sampled as biological replicates.

For the biomass calculation and estimation of total plant production, four-month-old tobacco plants were harvested at the end of the assay and their leaves classified according to leaf age; a correction factor based on production levels measured at different leaf ages was used to estimate total yields.

*N. benthamiana* leaf samples for intact tissue HSPME VOCs analysis were collected from young leaves ranging in weight from 150 to 350 mg and rolled inside the vials. Emission values were later normalized using leaf weight. Tobacco leaf samples for volatile release in dynamic condition assays were represented by young leaves from the upper part of the plant, 30-35 cm long.

Flower sampling was carried out at pre-anthesis (-1 day for *N. benthamiana*) and with completely open flowers, for both *N. benthamiana* and *N. tabacum* flowers.

### Analysis of volatile organic compounds (VOCs)

For powdered samples, 50 mg of frozen, ground leaf or flower tissue were weighed in a 10 mL or 20 mL headspace screw-cap vial and stabilized by adding 1 mL of 5M CaCl_2_ and 150 µL of 0.5 M EDTA, pH=7.5, after which they were immediately bath-sonicated for 5’. Volatile compounds were captured by means of headspace solid phase microextraction (HS-SPME) with a 65 µm polydimethylsiloxane/divinylbenzene (PDMS/DVB) SPME fiber (Supelco, Bellefonte, PA, USA). Volatile extraction was performed automatically by means of a CombiPAL autosampler (CTC Analytics, Zwingen, Switzerland).

For the T_0_, T_1_ and T_2_ generations in *N. benthamiana*, and for the T_0_ and T_1_ generations in *N. tabacum*, analyses were made using a PEGASUS 4D mass spectrometer (LECO Corporation, St. Joseph, MI, USA). Vials were first incubated at 50°C for 10’ under 500 rpm agitation. The fiber was then exposed to the headspace of the vial for 20’ under the same conditions of temperature and agitation. Desorption was performed at 250°C for 1’ (splitless mode) in the injection port of a 6890 N gas chromatograph (Agilent Technologies, Santa Clara, CA, USA) coupled to PEGASUS 4D mass spectrometer (LECO Corporation). After desorption, the fiber was cleaned in a SPME fiber conditioning station (CTC Analytics) at 250°C for 5’ under a helium flow. Chromatography was performed on a BPX-35 (30 m, 0.32 mm, 0.25 µm) capillary column (SGE) with helium as the carrier gas at a constant flow of 2 mL/min. The oven conditions started with an initial temperature of 40°C for 2’, 5°C/min ramp until 250°C, and a final hold at 250°C for 5’. Data was recorded in a PEGASUS 4D mass spectrometer (LECO Corporation) in the 35-300 m/z range at 20 scans/s, with electronic impact ionization at 70 eV. Chromatograms were processed by means of the ChromaTOF software (LECO Corporation).

For the T_3_ generation *N. benthamiana LPPS* and *CPPS* transformants, the T_2_ and T_3_ generation tobacco *LPPS* and *CPPS* transformants, and the T_0_ and T_1_ *LiLPPS LiAAT4* tobacco transformants, desorption was performed at 250°C for 1’ (splitless mode) in the injection port of a 6890 N gas chromatograph (Agilent Technologies) coupled to a 5975B mass spectrometer (Agilent Technologies). Chromatography was performed on a DB5ms (60 m, 0.25 mm, 1 µm) capillary column (J&W) with helium as the carrier gas at a constant flow of 1.2 mL/min. Oven conditions were the same indicated above. Data was recorded in 5975B mass spectrometer in the 35-300 m/z range at 20 scans/s, with electronic impact ionization at 70 eV. Chromatograms were processed by means of the Agilent MassHunter software (Agilent Technologies).

For intact tissue assays, leaf or flower tissue samples were weighed and placed in a 20 mL headspace screw-cap vial. Volatile compounds were captured by means of headspace solid phase microextraction (HS-SPME) with a 65 μm polydimethylsiloxane/divinylbenzene (PDMS/DVB) SPME fiber (Supelco). Volatile extraction was performed automatically by means of a CombiPAL autosampler (CTC Analytics). Vials were first incubated at 30°C for 10’ under 500 rpm agitation.

Chromatograms were processed by means of the Agilent MassHunter software. Identification of compounds was made by comparison of both retention time and mass spectrum with pure standards (lavandulol, chrysanthemol, linalool and lavandulyl acetate) and by comparison between mass spectrum for each compound with those of the NIST 2017 spectral library. Every compound quantification was corrected with the value of the daily deviation of a master mix, processed and analyzed every day. A linalool internal standard (IS) was always added as a control of hour drift.

The quantification of monoterpenoid compounds emitted either by whole plants, or by detached leaves or flowers was carried out by volatile collection in dynamic conditions. Individual plants for *N. benthamiana* or 50 g of tobacco leaves were placed inside 5 L glass reactors (25 cm high × 17:5 cm diameter flask) with a 10 cm open mouth and a ground glass flange to fit the cover with a clamp. The cover had a 29/32 neck on top to fit the head of a gas washing bottle and to connect a glass Pasteur pipette downstream to trap effluents in 400mg of Porapak-Q 80-100 (Waters Corporation, Milford, MA, USA) adsorbent. For tobacco flowers, around 3 g of flowers were placed inside 1.3 L glass chambers (50 cm length × 6 cm diameter cylinder). Plant and leaf samples were collected continuously for 72 h, and flower samples for 48 h, by using an ultrapurified-air stream, provided by an air compressor (Jun-air Intl. A/S, Norresundby, Denmark) coupled with an AZ 2020 air purifier system (Claind Srl, Lenno, Italy) to provide ultrapure air (amount of total hydrocarbons < 0:1 ppm). In front of each glass reactor, an ELL-FLOW digital flowmeter (Bronkhorst High-Tech BV, Ruurlo, The Netherlands) was fitted to provide an air push flow of 100 mL/min during sampling. Volatiles trapped in the Porapak Q cartridges were eluted with 3 mL pentane. Solvent extracts were concentrated under a gentle nitrogen stream up to 500 µL and 25 µL of an internal standard solution (100 μg/mL in dichloromethane) were added to the sample prior to the chromatographic analysis for quantification of the target molecules.

### Solvent extraction from plant tissues

The total quantity of pheromone compounds accumulated in each plant or leaf bunch was extracted with toluene (TLN). Plant samples (ca. 3 g), mixed with fine washed sand (1 : 1, plant : sand, w/w), were manually ground with a mortar to aid in tissue breakdown and facilitate the extraction. The resulting material was then transferred to 50 mL centrifuge tubes with 10 mL TLN. The extraction process was assisted by magnetic agitation for 12 h and finally by ultrasound in a Sonorex ultrasonic bath (Bandelin electronic, Berlin, Germany) for 30’. A 1 mL sample of the resulting extract was filtered through a PTFE syringe filter (0.25 μm). Twenty-five μL of an internal standard solution (100 μg/mL in dichloromethane) were added to the sample prior to the chromatographic analysis for quantification of the target molecules.

### Quantification of target compounds

The quantification was performed by gas chromatography coupled to mass spectrometry (GC-MS), using an internal standard. A straight chain fluorinated hydrocarbon ester (heptyl 4,4,5,5,6,6,7,7,8,8,9,9,9-tridecafluorononanoate; TFN) was selected as the internal standard to improve both sensitivity and selectivity for MS detection (Gavara et al. 2020).

One µL of each extract was injected in a Clarus 690 gas chromatograph (Perkin Elmer Inc., Wellesley, MA) coupled to a Clarus SQ8T MS instrument operating in full scan mode and using EI (70 eV). The GC was equipped with a ZB-5MS fused silica capillary column (30m × 0.25mm i.d. × 0.25 μm; Phenomenex Inc., Torrance, CA). The oven was held at 60°C for 1 min then was raised by 10°C/min up to 120°C, maintained for 4 min, raised by 10°C/min up to 130°C, and finally raised by 20°C/min up to 280°C held for 2 min. The carrier gas was helium at 1 mL/min. The GC injection port and transfer line were programmed at 250 °C, whereas the temperature of the ionization source was set at 200°C. Chromatograms and spectra were recorded with GC-MS Turbomass software version 6.1 (PerkinElmer Inc.).

The amount of each compound and the corresponding chromatographic areas were connected by fitting a linear regression model, *y* = a + b*x*, where *y* is the ratio between compound and TFN areas and *x* is the amount of compound.

### Chlorophyll Index measurements

For T_3_ generation *N. benthamiana* and *N. tabacum LPPS* and *CPPS* transformants, and T_1_ *NtLPPS-AAT4* tobacco transformants, chlorophyll index (C.I.) data were collected with a Dualex-A optical sensor (Dualex Scientific® (Force-A, Orsay, France). Three leaves per plant were sampled for each leaf stage: young and adult leaves in pre-flowering plants, and young, adult and early senescent leaves in post-flowering plants, using the same criteria specified for plant sampling.

### Synthesis of pure standards for GC-MS

#### Linalool

Standard sample of linalool was commercially acquired from Sigma-Aldrich.

#### Racemic Lavandulol

A solution of methyl acrylate (1.5 g, 0.013 mol) in dry tetrahydrofuran (5 ml) was slowly added to a cooled solution of lithium diisopropylamide at -40 °C (2.0 M in THF, 7.9 ml, 1.2 eq) under nitrogen. After 60 min of continuous stirring, prenyl bromide (2.22 g 0.014 mol, 1.15 eq) were added, and the solution was warmed up to room temperature and stirred for an additional 5 h. The reaction was quenched by the addition of saturated ammonium chloride solution (5 ml) and extracted with Et_2_O (3 X 15 ml). The combined organic phases were successively washed with HCl 1M (1 X 5 ml), NaHCO_3_ 10 % (1X 5 ml) and brine (1 X 10 ml), dried with anhydrous magnesium sulphate, and the solvent evaporated under vacuum. The crude material was dissolved in anhydrous tetrahydrofurane (4 ml) and slowly added to a suspension of LiAlH_4_ (0.67 g, 0.017 mol, 1.35 eq.) in dry THF (5 ml) at 0 °C under argon atmosphere. After 3 h of continuous stirring, sodium sulphate decahydrated (Glauber’s salt) was carefully added to the suspension until a clear solid was formed (hydrogen formed during the quenching was removed with a continuous stream of nitrogen). The solid was filtered off through a celite pad, and the solvent evaporated under vacuum. The crude material was purified by column chromatography using a mixture of hexane:acetate (8:2) as eluent. Evaporation of the solvent of the corresponding fractions afforded pure lavandulol (1.4 g, 96 % purity by GC-FID, 70 % overall yield for two steps). The spectroscopical properties of lavandulol were fully coincident with those described in the literature (Pepper *et al*. 2014).

#### Lavandulyl acetate

Triethyl amine (1.05 ml, 7.6 mmol, 1.8 eq.) and acetic anhydride (0.46 ml, 4.8 mmol, 1.15 eq.) were subsequently added to a solution of lavandulol (0.65 g, 4.2 mmol) in dry dichloromethane (8 ml) at room temperature. After 5 h of continuous stirring, the reaction was poured in dichloromethane (15 ml) and subsequently washed with HCl 1M (1 X 10 ml), NaHCO_3_ 10 % (1X 10 ml) and brine (1 X 10 ml), dried with anhydrous magnesium sulphate, and the solvent evaporated under vacuum. The crude material was purified by column chromatography using a mixture of hexane:acetate (9:1) as eluent. Evaporation of the solvent of the corresponding fractions afforded pure lavandulol (0.78 g, 98 % purity by GC-FID, 95 % yield). The spectroscopical properties of lavandulol acetate were fully coincident with previously reported data (Cross *et al*. 2004).

#### Chrysantemol

A solution of chrysanthemic acid (mixture of isomers, 3 g, 18 mmol) in anhydrous tetrahydrofurane (6 ml) was slowly added to a suspension of LiAlH_4_ (1.4 g, 38 mmol, 2 eq.) in dry THF (20 ml) at 0 °C under argon atmosphere. The reaction mixture was warm up to room temperature and, after 5 h of continuous stirring, sodium sulphate heptahydrated (Glauber’s salt) was carefully added to the suspension until a clear solid was formed (hydrogen formed during the quenching was removed with a continuous stream of nitrogen). The solid was filtered off through a celite pad, and the solvent evaporated under vacuum. The crude material was purified by column chromatography using a mixture of hexane:acetate (7:3) as eluent. Evaporation of the solvent of the corresponding fractions afforded pure (+)-*trans*-chrysanthemol (2.3 g, 96 % purity by GC-FID, 85 % yield). The spectroscopical properties of (+)-*trans*-chrysanthemol were fully coincident with those described in the literature (Dufour et al. 2012).

### Statistical analysis

Statistical analyses were performed using the Past4 software (Hammer *et al*., 2001) and GraphPad Prism v8.0.2 (GraphPad Software, San Diego, CA, USA).

## Results

### 1. Volatile irregular monoterpenoid alcohols lavandulol and chrysanthemol are efficiently produced in tobacco and *N. benthamiana*

The production of irregular monoterpenoids from their DMAPP precursor (Figure 1A) in non-specialized leaf cells was first assayed by transient agroinfiltration in *N. benthamiana* of the genes encoding the corresponding isoprenyl transferases. The coding sequences of *Li*LPPS and *Tc*CPPS were assembled in separate GoldenBraid vectors, each of them regulated by the Cauliflower Mosaic Virus 35S promoter (pCaMV35S) and the nopaline synthase terminator (tNOS) (Figure 1B,C). The volatile organic compound (VOC) composition of infiltrated leaves was analyzed 5 days post-infiltration (dpi) by headspace solid-phase micro-extraction (HSPME) gas chromatography/mass spectrometry (GC/MS). Production of the volatile monoterpenoid alcohols lavandulol and chrysanthemol was successfully detected in *LiLPPS*- and *TcCPPS*-agroinfiltrated samples, respectively (Figure 1B,C). Relative quantifications are shown in Figure 1E. None of these products was detectable in negative controls (Figure 1D,E). In *TcCPPS*-infiltrated leaves other related volatile monoterpenoids were detected, namely Artemisia and Yomogi alcohols and santolinatriene, as identified by the NIST mass spectral library (2017). Both Artemisia and Yomogi alcohol are known to derive from the chrysanthemyl cation in aqueous environments, through the rupture of its C(1’)-C(3’) cyclopropane bond (Poulter *et al*., 1977; Rivera *et al*., 2001).

Given the success obtained in transient expression, the generation of stable lines producing lavandulol and chrysanthemol was attempted both for *N. benthamiana* and tobacco. Twelve T_0_ *LiLPPS N. benthamiana* (*NbLPPS*) plants and four T_0_ *LiLPPS* tobacco (*NtLPPS*) plants were recovered on selective media. Similarly, seven *TcCPPS N. benthamiana* (*NbCPPS*) and three *TcCPPS* tobacco (*NtCPPS*) T_0_ plants were obtained. The production of targeted monterpenoids in all four transformation experiments was followed for individual plants in T_0_, T_1_ and T_2_ generations, and the results are shown in Figure S1. Selection for best-performing lines through generations was made based on production/growth balance, and general trends remained stable up to the T_2_ generation. *NtLPPS* plants (especially line *NtLPPS*_3_1) were more productive than their *N. benthamiana* counterparts, with up to 7-fold the lavandulol levels detected in *NbLPPS* plants, except for line *NbLPPS*_11_2_4, whose levels reached 50% those of the best tobacco producers. *NtCPPS* plants produced over 4 times more chrysanthemol than their *NbCPPS* counterparts. In both species, plants producing lavandulol showed yellowing and slower growth, in contrast to non-producer transgenic plants, which were comparable to WTs, and a similar trend was observed for *NbCPPS* and *NtCPPS* plants. Especially for *NbCPPS* and *NtCPPS*, production was associated with premature blossom drop and frequent failure to reach fruit set (not shown). In all chrysanthemol-producing plants, the derived products Artemisia and Yomogi alcohols and santolinatriene were detected, most likely being breakdown products generated during sample processing (Figure S2). This points to a likely under-estimation of chrysanthemol in these samples. However, while taking this factor into consideration, at this point we used these assays only as relative quantification methods to compare different plants within the *NbCPPS* and *NtCPPS* populations.

### 2. Lavandulol and chrysanthemol production is higher in the earlier developmental stages and in young leaves of *N. benthamiana* and tobacco transgenic lines

For a thorough characterization of transgenic *N. benthamiana* and tobacco plants, we analyzed the levels of the target compounds in leaves at different developmental stages of the plant, as well as at different stages of leaf development, in a uniform T_3_ generation. Two tobacco (*NtLPPS*_1_3_2 and *NtLPPS*_3_1_3) and two *N. benthamiana* (*NbLPPS*_11_2_4 and *NbLPPS*_5_2_2) lines were selected based on their lavandulol production levels. For *NbCPPS* and *NtCPPS*, one vigorous line with a single T-DNA insertion, characterized by high production rates, was selected for each *Nicotiana* species. Overexpression of monoterpenes is known to affect plant growth and fertility, a trend we had observed in the T_1_ and T_2_ generations of plants producing lavandulol and chrysanthemol. As expected, all T_3_ transgenic lines showed a delay in flowering time compared to WTs (Figure 2A,E). This delay consists of a 20-30% increase in the number of days in soil before flowering in *N. benthamiana*, while in tobacco the delay can be greater, reaching 93% in *NtLPPS*_3_1_3. No significant differences between genotypes are found regarding plant height except for *NtLPPS*_3_1_3 (Figure 2B,F), but a reduction in biomass is observed. In *N. benthamiana*, both *NbLPPS* lines show a significant reduction in biomass (estimated as FW after 100 days in soil) which can range between 60 and 80% compared to both WT and *NbCPPS* (Figure 2C,D). In tobacco, differences in size and plant biomass are not significant between WT and *NtCPPS*, but they are for *NtLPPS*, whose biomass is reduced between 35 and 75% (Figure 2G,H). The chlorophyll index (C.I.) is another parameter for measuring loss of fitness due to overexpression of monoterpenoids, since these compounds often induce chlorosis. In *N. benthamiana*, no significant differences are found in the C.I. of *NbCPPS* compared to WTs, while for *NbLPPS* there is a reduction at both the pre- and post-flowering stages (Figure S3). In tobacco, both *NtLPPS* and *NtCPPS* show a reduction in C.I. at the pre-flowering stage, which disappears post-flowering (Figure S3). In all cases, differences correlate with higher monoterpenoid production (see also Figure 3). No correlation was found between the number of copies of the transgene and production levels or other phenotypic effects. According to segregation patterns, *NbLPPS*_11_2_4 and *NtLPPS*_1_3_2 had a single T-DNA insertion, while *NbLPPS*_5_2_2 and *NtLPPS*_3_1_3 carried multiple transgene copies. While plants of the multiple-copy line *NtLPPS*_3_1_3 show the most severe pleiotropic effects, in *N. benthamiana* the single-copy line *NbLPPS*_11_2_4 has the greatest yields and most severe phenotypes.

**Figure 2.**
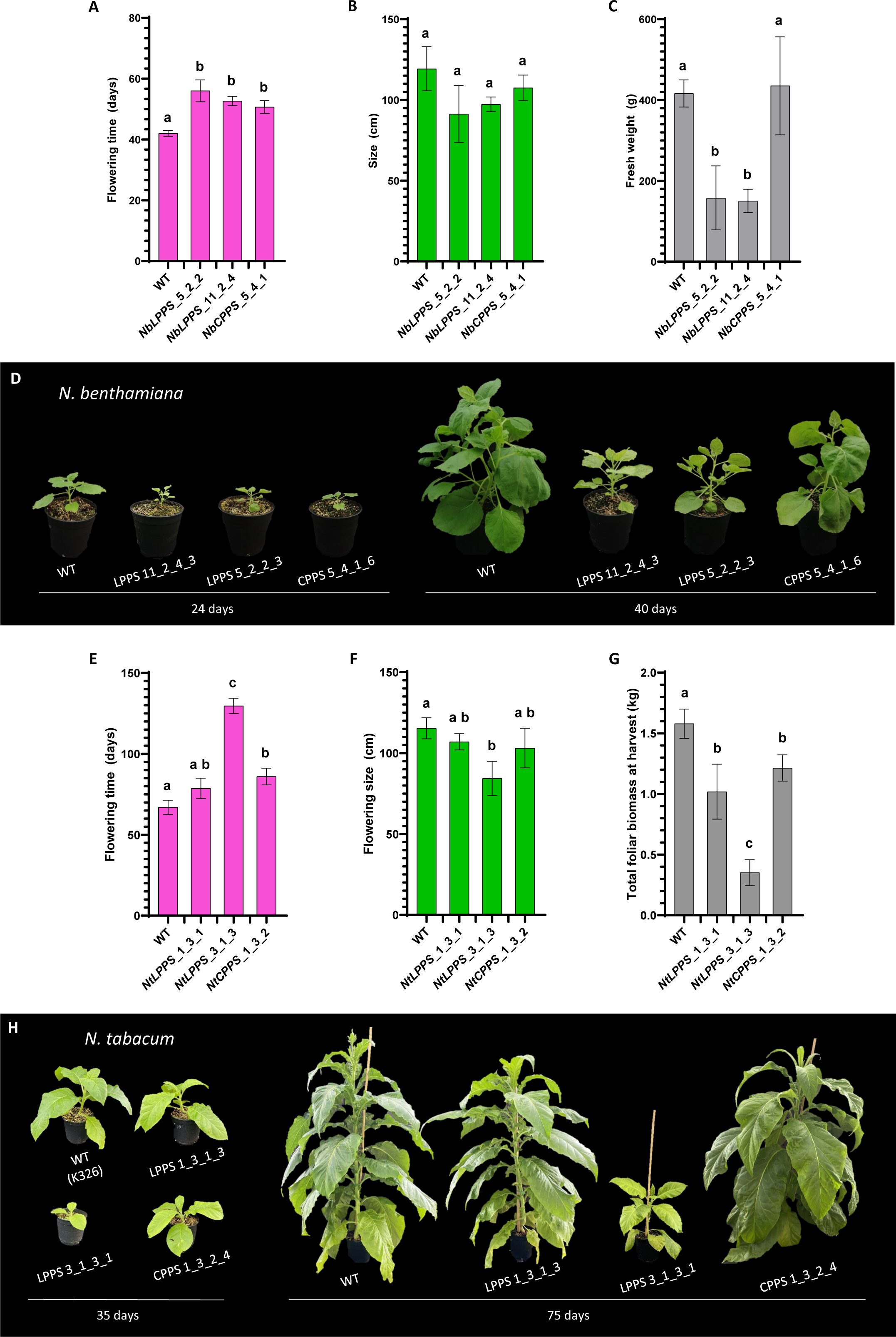
Physiological effect of irregular monoterpene production in T_3_ N. benthamiana and tobacco plants. Comparison of flowering time **(A)**, plant size at 100 days **(B)** and total biomass reached at 100 days **(C)** in *NbLPPS*, *NbCPPS* and WT plants. In panel (A), time measured as days from transfer to soil to anthesis of the first flower. **(D)** Phenotype of *NbLPPS* and *NbCPPS* plants compared to WT at 24 and 48 days in soil. Comparison of flowering time **(E)**, plant size at flowering **(F)** and total foliar biomass accumulated at harvest time (140 days) **(G)** in *NtLPPS*, *NtCPPS* and WT plants. In panel **(E)**, time measured as days from transfer to soil to formation of the floral meristem. **(H)** Phenotype of *NtLPPS* and *NtCPPS* plants compared to WT at 35 and 75 days in soil. Values are the mean and standard deviation of at least 3 independent plants of each line. Error bars with the same letter are not significantly different (one-way ANOVA with post-hoc Tukey HSD at the 5% level of significance).

**Figure 3.**
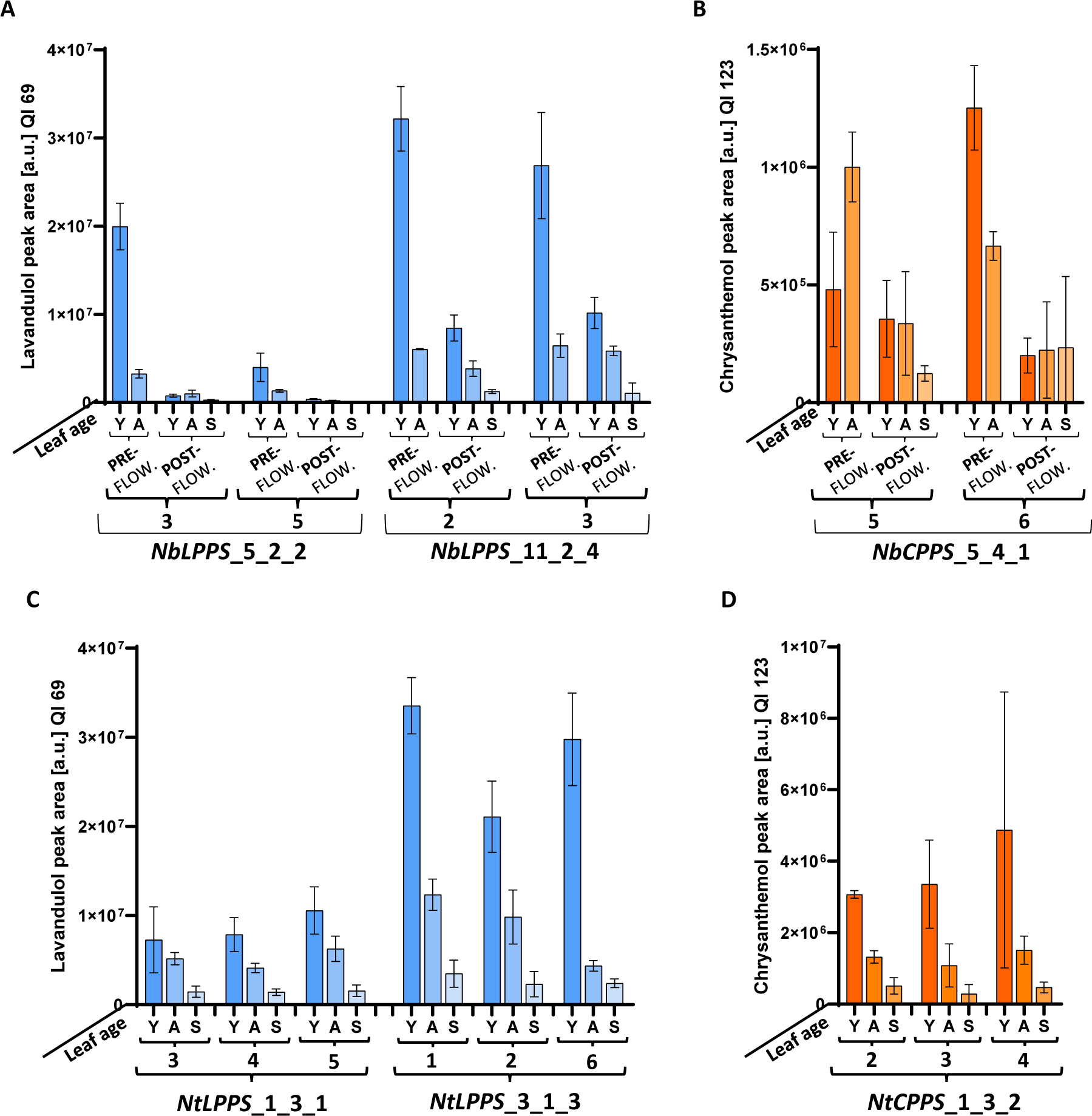
Comparison of lavandulol and chrysanthemol production in leaves at different developmental stages of T_3_ *N. benthamiana* and tobacco plants. (A) Lavandulol production in *NbLPPS*_11_2_4 and *NbLPPS*_5_2_2 lines. **(B)** Chrysanthemol production in *NbCPPS*_5_4_1 line. **(C)** Lavandulol production in *NtLPPS*_1_3_1 and *NtLPPS*_3_1_3 lines. **(D)** Chrysanthemol production in *NtCPPS*_1_3_2 line. In panels (A) and (B) values are reported for two plant growth stages (pre-flowering and post-flowering) and for three leaf developmental stages (Y=young, A=adult, S=senescent). In panels (C) and (D) values are reported for three types of leaves (Y=young, A=adult, S=senescent). Production values represent the mean and standard deviation of n = 3 biological replicates (independent leaves). All data for this figure, together with statistical analyses (two-way ANOVA) are reported in Supplementary Data File S1.

The monoterpenoid content of leaves was assessed pre- and post-anthesis in *N. benthamiana* plants. Pre-anthesis, two types of leaves were analyzed (Y=young and A=adult), while post-anthesis a third type of leaf (S=senescent) was included. The analysis of the monoterpenoid content in different individuals and tissues (Figure 3A,B and Data File S1) shows that the factor affecting productivity the most is the type of tissue. Plants within each line show homogeneous phenotypes in terms of production levels and fitness, and no significant differences are found between plants descending from the same T_2_ parental. Overall, young leaves of pre-anthesis plants are the most productive vegetative tissues in both *NbLPPS* and *NbCPPS* lines, followed by the adult leaves of these pre-anthesis plants (Figure 3A,B). We observed a decrease in the levels of the target monoterpenoids in fully flowering plants, diminishing with the increase in leaf age (Figure 3A,B). In *NbLPPS* and *NbCPPS*, adult leaves of pre-flowering plants and young leaves of post-anthesis plants usually show similar production levels. Senescent leaves are still productive, even if a clear decline was observed for all analyzed compounds. In tobacco, only one sampling was performed just prior to anthesis, and three leaf types were collected (Y=young, A=adult and S=senescent). The same decrease in monoterpenoid production with increasing leaf age was identified in *NtLPPS* and *NtCPPS* plants: young leaves stand out as the most productive vegetative tissue in tobacco (Figure 3C,D and Data File S1). *NbLPPS*_11_2_4 and *NtLPPS*_3_1_3 have comparable levels of lavandulol, especially in young leaves. By contrast, tobacco produced more chrysanthemol than *N. benthamiana*, with young leaves of *NtCPPS*_1_3_2 producing more than twice the levels detected in the best performing *NbCPPS*_5_4_1 young pre-flowering leaves.

For a more accurate measure of monoterpenoid production, solvent extraction was used to obtain absolute quantifications; for this, materials from the best-yielding conditions and single copy lines were analyzed. From *NbLPPS*_11_2_4 whole young T_3_ plants, almost 35 µg/g FW of lavandulol were obtained, in contrast to 0.6 µg/g FW of chrysanthemol in *NbCPPS*_5_4_1 plants (Table 1). These quantifications correlate with the accumulation of lavandulol and chrysanthemol observed in relative quantifications (Figure 3A,B). A similar trend was observed in tobacco: an average of 22.55 µg/g FW of lavandulol were retrieved from young leaves of *NtLPPS*_1_3_1 plants, while extraction from young leaves of *NtCPPS*_1_3_2 plants yielded an average of 0.6 µg/g FW of chrysanthemol (Figure 3C,D and Table 1).

**Table 1.**
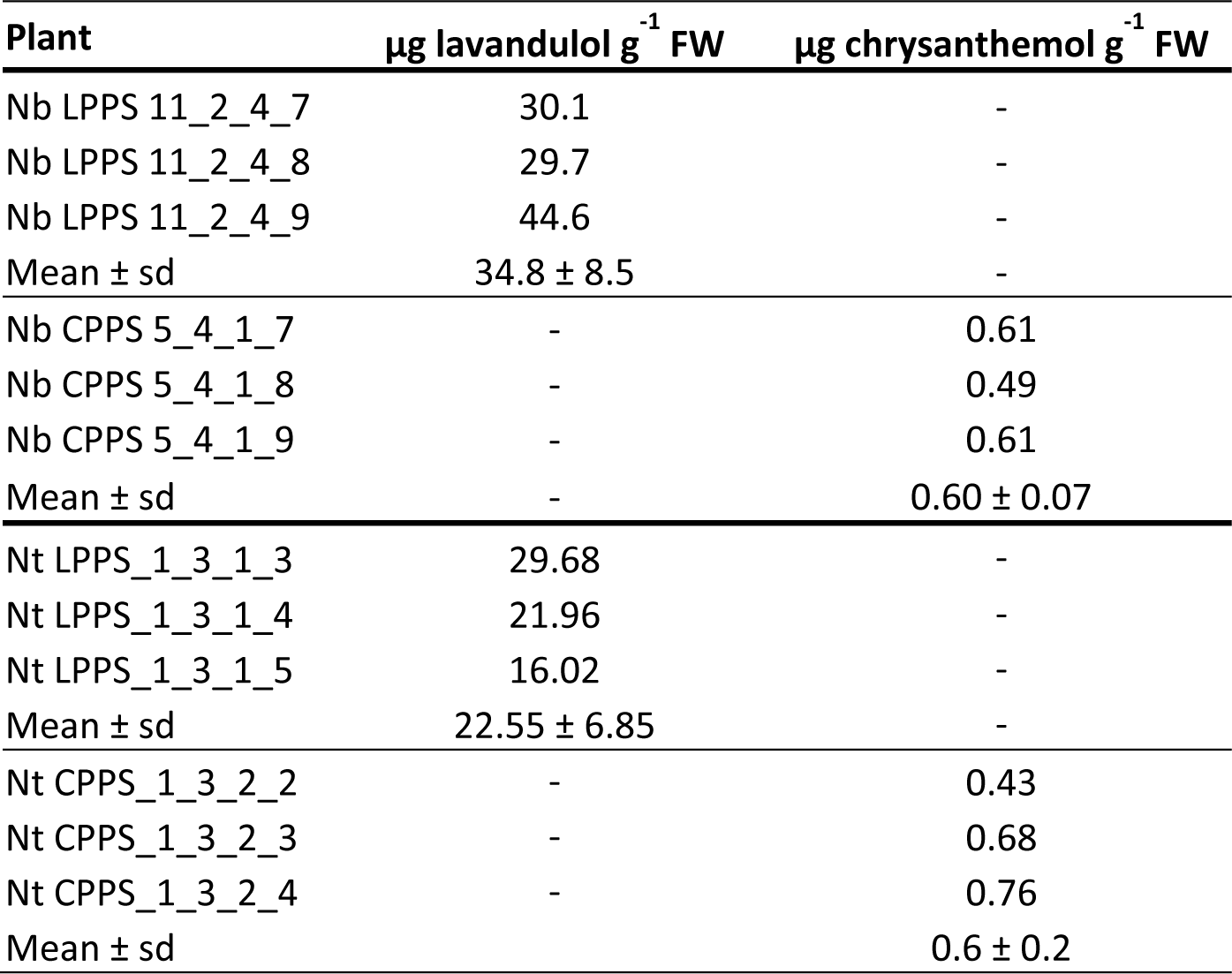
Quantity (µg) of lavandulol and chrysanthemol obtained from young whole T_3_ *NbLPPS* and *NbCPPS* plants and from leaves of young T_3_ *NtLPPS* and *NtCPPS* plants by solvent extraction and GC/MS/MS quantification.

| Plant | $\mu\text{g lavandulol g}^{-1} \text{ FW}$ | $\mu\text{g chrysanthemol g}^{-1} \text{ FW}$ |
| --- | --- | --- |
| Nb LPPS 11_2_4_7 | 30.1 | - |
| Nb LPPS 11_2_4_8 | 29.7 | - |
| Nb LPPS 11_2_4_9 | 44.6 | - |
| Mean ± sd | $34.8 \pm 8.5$ | - |
| Nb CPPS 5_4_1_7 | - | 0.61 |
| Nb CPPS 5_4_1_8 | - | 0.49 |
| Nb CPPS 5_4_1_9 | - | 0.61 |
| Mean ± sd | - | $0.60 \pm 0.07$ |
| Nt LPPS_1_3_1_3 | 29.68 | - |
| Nt LPPS_1_3_1_4 | 21.96 | - |
| Nt LPPS_1_3_1_5 | 16.02 | - |
| Mean ± sd | $22.55 \pm 6.85$ | - |
| Nt CPPS_1_3_2_2 | - | 0.43 |
| Nt CPPS_1_3_2_3 | - | 0.68 |
| Nt CPPS_1_3_2_4 | - | 0.76 |
| Mean ± sd | - | $0.6 \pm 0.2$ |

### 3. The potential for volatilization of lavandulol and chrysanthemol in vegetative and reproductive tissues of transgenic lines

Volatilization from stabilized ground tissue in HSPME vials was adopted as a high-throughput screening method to estimate the content of VOCs, requiring minimal amounts of manipulation and reagents, which allowed us to compare a great number of samples in parallel, while also getting a general picture of the tissue volatilome beyond extraction of single classes of compounds. Other approaches (in addition to solvent extraction) were deemed necessary to characterize the production and emission of monoterpenoid alcohols more accurately, keeping in mind the prospective use of plants as live bio-dispensers. Two strategies were followed: *i*) incubating intact samples in HSPME vials and analyzing the emitted compounds and *ii*) analyzing volatiles emitted by intact plants or leaves under dynamic conditions. Flowers emit great quantities of volatiles in many plant species (Loughrin *et al*., 1991; Dudareva *et al*., 2013; Adebesin *et al*., 2017): we wondered if *N. benthamiana* or tobacco flowers could be responsible for a considerable percentage of the total volatile monoterpenoid production of our transgenic plants. In addition to leaves, both intact and ground flowers from transgenic plants were analyzed by HSPME GC/MS.

For *NbLPPS*, the lavandulol emitted by intact leaves represented around one eighth of the lavandulol measured in homogenized samples (Figure 4A). The lavandulol detected in ground *NbLPPS* flowers is 22% of that of ground leaves, but the volatilized portion is higher in flowers, with the emitted fraction reaching almost 30% of that measured in ground flower samples (Figure 4A). Lavandulol emission was similar when intact flowers and leaves were compared. An analogous trend was observed for *NbCPPS*: chrysanthemol emission was almost undetectable in intact leaves, while being considerably higher in ground tissue (Figure 4B), and it was also detected in intact flowers, which emit around 90% more chrysanthemol than leaves per biomass unit. The production of lavandulol and chrysanthemol did not depend on the developmental stage of *N. benthamiana* flowers, since no differences were found between flowers before and after anthesis for either compound (Figure S4). When quantifying emission from young intact *N. benthamiana* plants under dynamic conditions, 160 ng/g FW/day lavandulol were detected in *NbLPPS* plants, four times the amount of chrysanthemol emitted by *NbCPPS* plants (40 ng/g FW/day, Table 2).

**Figure 4.**
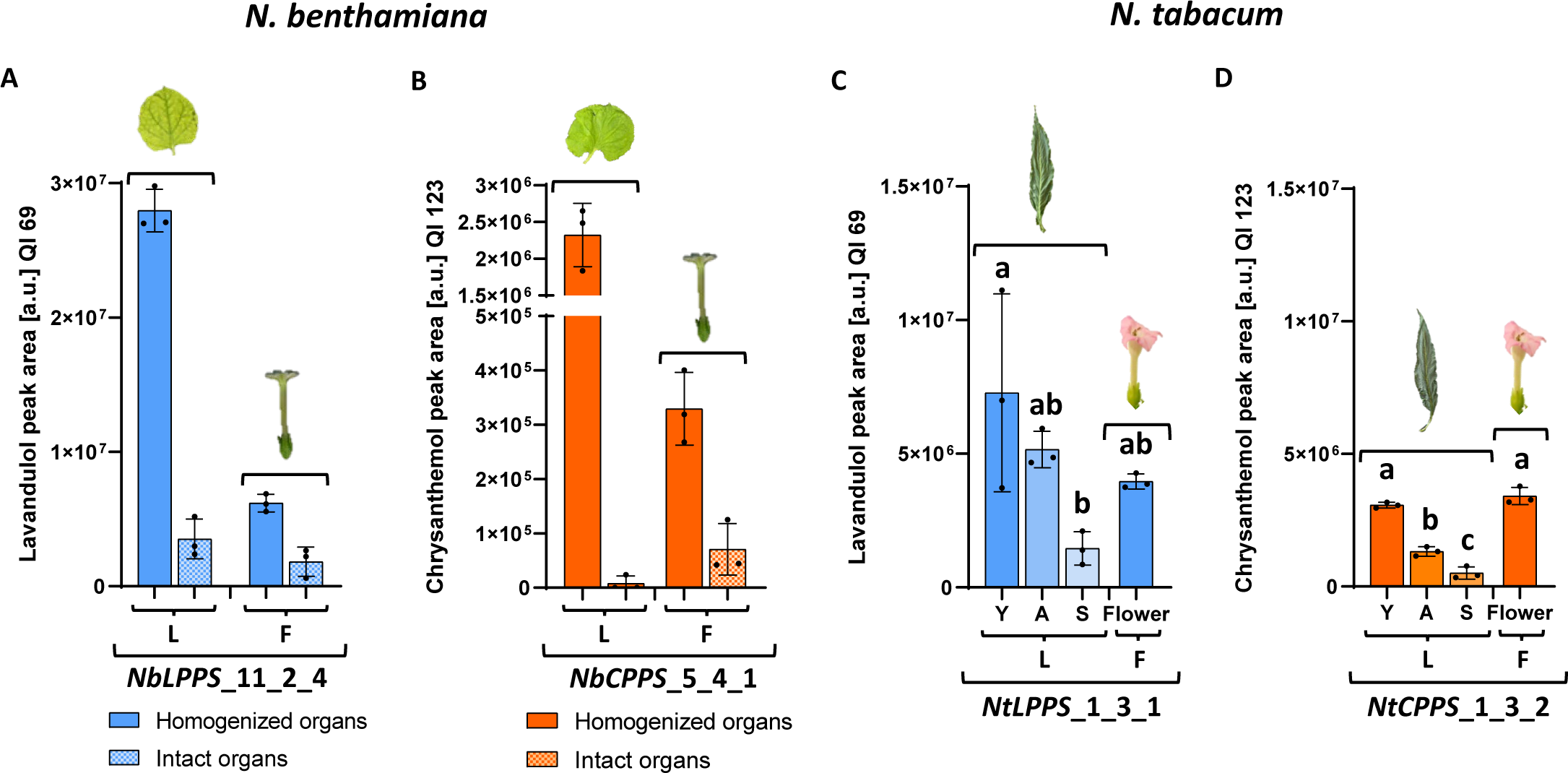
Production of lavandulol and chrysanthemol in T_3_ *N. benthamiana* and tobacco plants in different tissue types. **(A)** Lavandulol production in vegetative and reproductive tissue in line *NbLPPS*_11_2_4, measured in homogenized and intact organs. **(B)** Chrysanthemol production in reproductive and vegetative tissue in line *NbCPPS*_5_4_1, measured in homogenized and intact organs. **(C)** Lavandulol production in vegetative and reproductive tissues in line *NtLPPS*_1_3_1, measured in homogenized leaves and flowers. **(D)** Chrysanthemol production in vegetative and reproductive tissue in line *NtCPPS*_1_3_2, measured in homogenized leaves and flowers. In all panels, leaf tissues are denoted as “L” and flower tissues as “F”. In panels (C) and (D) values are reported for three types of leaves (Y=young, A=adult, S=senescent). Values represent the mean and SD of n=3 biological replicates (independent organs). Comparisons (Student’s *t* and Mann-Whitney’s test for *N. benthamiana* and one-way ANOVA with HSD Tukey’s posthoc test, p<0.05) were carried out between samples undergoing the same treatment (homogeneized or intact). Letters identify significance groups.

**Table 2.** Quantity (ng) of lavandulol and chrysanthemol released by whole T_3_ *NbLPPS* and *NbCPPS* young individuals, and by T_3_ *NtLPPS* and *NtCPPS* young leaves and flowers, measured by volatile collection and GC/MS/MS quantification.

| Plant | Material | ng lavandulol g<sup>-1</sup> FW day<sup>-1</sup> | ng chrysanthemol g<sup>-1</sup> FW day<sup>-1</sup> |
| --- | --- | --- | --- |
| Nb LPPS 11_2_4_7 | Full plant (young) | 218.42 | - |
| Nb LPPS 11_2_4_8 | Full plant (young) | 96.33 | - |
| Nb LPPS 11_2_4_9 | Full plant (young) | 159.91 | - |
| Mean ± sd | Full plant (young) | 158.22 ± 61.06 | - |
| Nb CPPS 5_4_1_7 | Full plant (young) | - | 62.95 |
| Nb CPPS 5_4_1_8 | Full plant (young) | - | 24.32 |
| Nb CPPS 5_4_1_9 | Full plant (young) | - | 44.98 |
| Mean ± sd | Full plant (young) | - | 44.08 ± 19.33 |
| Nt LPPS_1_3_1_3 | Young leaves | 48.61 | - |
| Nt LPPS_1_3_1_4 | Young leaves | 54.59 | - |
| Nt LPPS_1_3_1_5 | Young leaves | 58.48 | - |
| Mean ± sd | Young leaves | 53.89 ± 4.97 | - |
| Nt LPPS_1_3_1_3 | Flowers | 168.21 | - |
| Nt LPPS_1_3_1_4 | Flowers | 348.7 | - |
| Nt LPPS_1_3_1_5 | Flowers | 392.11 | - |
| Mean ± sd | Flowers | 303.01 ± 118.74 | - |
| Nt CPPS_1_3_2_2 | Young leaves | - | 19.04 |
| Nt CPPS_1_3_2_3 | Young leaves | - | 21.72 |
| Nt CPPS_1_3_2_4 | Young leaves | - | 13.54 |
| Mean ± sd | Young leaves | - | 18.1 ± 4.17 |
| Nt CPPS_1_3_2_2 | Flowers | - | 45.49 |
| Nt CPPS_1_3_2_3 | Flowers | - | 36.92 |
| Nt CPPS_1_3_2_4 | Flowers | - | 14.08 |
| Mean ± sd | Flowers | - | 32.16 ± 16.24 |

In tobacco transgenic lines, lavandulol and chrysanthemol are produced at comparable levels in ground flowers and young leaves (Figure 4C,D). In terms of volatility, flowers emit 5.6 times more lavandulol than leaves per biomass unit, while no significant differences are found for chrysanthemol between the two tissues (Table 2). Comparing emission of lavandulol and chrysanthemol, under dynamic conditions the lavandulol released by *NtLPPS* young leaves (53.89 ng/g FW/day) is almost three times more abundant than the chrysanthemol released by *NtCPPS* leaves (18.1 ng/g FW/day). Again, flowers proved to be better emitters than leaves: *NtLPPS* flowers released almost six-fold more lavandulol than vegetative tissues of the same biomass, and *NtCPPS* flowers almost doubled the chrysanthemol emission of the leaves (Table 2).

Comparing the two biofactories, monoterpenoid alcohol levels are similar in ground floral tissues, but the potential for emission is greater in tobacco than in *N. benthamiana* flowers. Compared to *NbLPPS*, *NtLPPS* leaves emitted one third of the lavandulol volatilized by *NbLPPS* whole plants; similarly, *NtCPPS* leaves emitted around half of the chrysanthemol released by *NbCPPS* whole plants, per biomass unit (Table 2). The two sets of values are not fully comparable (whole plants *vs*. detached leaves), yet we can hypothesize that, considering the production levels estimated for the different kind of leaves, the greater biomass of adult tobacco plants would allow a considerably greater total emission. Interestingly, the chrysanthemol-derived compounds Artemisia and Yomogi alcohols found in ground samples are barely detectable, if at all, in the emitted volatilome of intact samples (Data File S2).

### 4. Lavandulol can be esterified to lavandulyl acetate by the *Li*AAT-4 acetyltransferase in tobacco

We wanted to test the ability of our plants to produce lavandulyl acetate as an example of an active pheromone compound derived from one of the assayed precursors. In addition to its value as a fragrance, (R)-lavandulyl acetate is a semiochemical found in the aggregation pheromone of the thrips *F. occidentalis* (Hamilton *et al*., 2005) and in the sex pheromone of the mealybug *D. grassii* (De Alfonso *et al*., 2012). Notably, Govindarajan & Benelli (2016) also highlighted its effectiveness as a mosquito larvicide with a LC50 of around 4 µg/ml in aqueous solution. The AAT4 acetyltransferase from *L. intermedia* (*Li*AAT4) acetylates monoterpenoid alcohols, including lavandulol (Sarker & Mahmoud, 2015). We first tested *LiAAT4* in *N. benthamiana* leaves: a construct containing the P35S::*LiAAT4*::tNOS transcriptional unit was agroinfiltrated on T_3_ *NbLPPS* plants (Figure 5A) and high levels of acetylation of the lavandulol substrate were obtained (Figure 5B), with an average 70% of the substrate detected in the absence of *Li*AAT4 converted to lavandulyl acetate.

**Figure 5.**
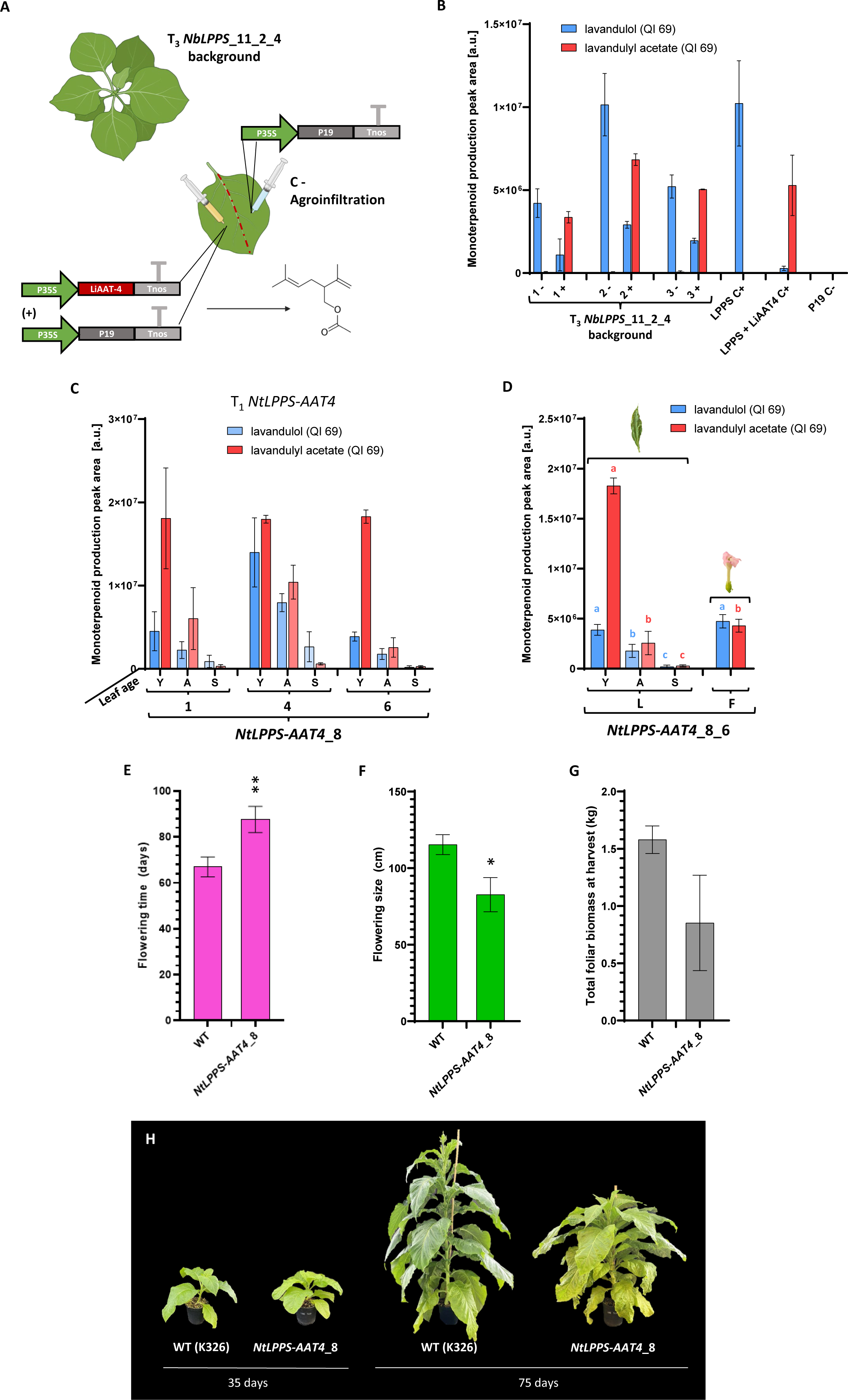
Esterification of lavandulol to lavandulyl acetate. **(A)** Constructs carrying the *LiAAT4* transgene controlled by the CAMV35s promoter and NOS terminator and the silencing suppressor P19, and the design of the agroinfiltration assay. **(B)** Levels of lavandulyl acetate obtained by infiltrating *Li*AAT4 in *NbLPPS* T_3_ plants (*NbLPPS*_11_2_4 background); for each individual, ‘-’ indicates infiltration with P19 alone, while ‘+’ indicates infiltration with *Li*AAT4 and P19. Positive controls are represented by WT leaves infiltrated with *Li*LPPS alone (LPPS C+) and with *Li*LPPS and *Li*AAT4 (LPPS AAT4 C+), always combined with P19. The negative control is represented by WT plants infiltrated only with P19 (C-). **(C)** Production of lavandulol and lavandulyl acetate in the T_1_ progeny of *NtLPPS_AAT4*_8; values are reported for three types of leaves (Y=young, A=adult, S=senescent). **(D)** Production of lavandulol and lavandulyl acetate in reproductive (indicated as F) and vegetative (indicated as L) tissues (ground samples) of T_1_ *NtLPPS_AAT4*_8 plants. Three types of leaves are analyzed (Y=young, A=adult, S=senescent). Comparison of flowering time **(E)**, plant size at flowering **(F)** and total foliar biomass accumulated at harvest time (140 days) **(G)** in *NtLPPS_AAT4* and WT plants. In panel (E), time measured as days from transfer to soil to formation of the floral meristem in T_1_ *NtLPPS_AAT4* and WT plants. **(H)** Phenotype of T_1_ *NtLPPS_AAT4*_8 plants compared to the WT at 35 and 75 days in soil. In panels (B), (C) and (D) values are the mean and standard deviation of 3 independent samples. In panels (E), (F) and (G), values are the mean and standard deviation of at least 3 independent plants of each line. *P*-values were calculated using Student’s *t*-test; \**P* ≤ 0.05, \*\**P* ≤ 0.01. The figure includes images from Biorender (biorender.com).

Given the greater biomass and general robustness of tobacco compared to *N. benthamiana*, we used the T_1_ tobacco line producing the highest levels of lavandulol, *NtLPPS*_3_1, for stable transformation with *LiAAT4*. In the *NtLPPS_AAT4* T_0_ population, different efficiencies were observed for the conversion of lavandulol to lavandulyl acetate, measured by HSPME GC-MS (Figure S5). Some individuals showed especially good acetylation rates, producing levels of lavandulyl acetate comparable to those of lavandulol in their parental line, with accordingly lower levels of lavandulol. However, a strong negative correlation was observed between production levels and fitness, and the plants with the highest lavandulyl acetate production were not able to produce viable seeds. For phenotypic characterization, the progeny of the plant with the highest production levels which allowed to collect viable seeds (*NtLPPS_AAT4*_8) was chosen. As for *NtLPPS* plants, we evaluated accumulation in different plant tissues. Like in T_3_ tobacco and *N. benthamiana* plants producing lavandulol and chrysanthemol, young leaves of T_1_ *NtLPPS-AAT4* plants were the most productive tissues (between 2- and 7-fold more than adult leaves), while the lowest levels of lavandulyl acetate were observed in senescent leaves, paralleling lavandulol availability (Figure 5C). Again, the greatest source of variation was the type of tissue. No significant differences were found between the levels of lavandulyl acetate in the three tested T_1_ individuals within each leaf type, while *NtLPPS-AAT4*_8_4 accumulated more alcohol in young and adult leaves than the other T_1_ plants. Considering the total monoterpenoid content of the three individuals, no significant differences were found. The same analyses performed for *NtLPPS* and *NtCPPS* were carried out on *NtLPPS-AAT4* to determine production and emission in flowers and leaves and to quantify lavandulyl acetate by solvent extraction and under dynamic conditions. In absolute quantifications following solvent extraction, lavandulyl acetate appeared to be less accumulated in leaf tissues than lavandulol (Table 3). These data contrast sharply with the results of the analyses of homogenized tissue, in which the acetate:alcohol ratio is always higher and favors the acetate (Figure 5C), making us wonder whether part of it might be lost during the extraction procedure. As for *in vivo* release from leaves under dynamic conditions, volatilization is higher for lavandulyl acetate than for lavandulol: the highest producing plants release on average 626.84 ng/g FW/day of lavandulyl acetate and 72.45 ng/g FW/day of lavandulol (Table 4), compared to 53.89 ng/g FW/day of lavandulol released by the *NtLPPS*_1_3_1 parental line (Table 2). Regarding flowers, in homogenized samples lavandulol and lavandulyl acetate levels are remarkably similar (Figure 5D), also being comparable to those of lavandulol in *NtLPPS* flowers (Figure 4C). Volatilization from intact flowers favors lavandulyl acetate over lavandulol, although in absolute terms the emission rate from flowers is on average 2.5-fold less abundant than it is from young leaves (253 vs 627 ng/gFW/day, see Table 4). The heterologous production of lavandulyl acetate, too, was accompanied by pleiotropic effects such as yellowing leaves, a delay in flowering and a lower biomass at flowering time (Figure 5E,F,G,H and Figure S3). Interestingly, when compared to the T_3_ tobacco plants derived from the same parental used for stacking the *LiAAT4* transgene, the phenotype of the plants producing only lavandulol (Figure 2H) was overall more severe than that of those producing high levels of lavandulyl acetate: *NtLPPS*_3_1_3 plants accumulate 22% of the leaf biomass of WT tobacco, while *NtLPPS_AAT4*_8 T_1_ plants reach 53%. However, the greatest reduction in chlorophyll index was found in *NtLPPS-AAT4* plants (Figure S3).

**Table 3.** Quantity (µg) of lavandulol and lavandulyl acetate obtained from young leaves of T_1_ *NtLPPS-AAT4* plants by solvent extraction and GC/MS/MS quantification.

| Plant | µg lavandulol g<sup>-1</sup> FW | µg lavandulyl acetate g<sup>-1</sup> FW |
| --- | --- | --- |
| Nt LPPS-AAT4_4_4 | 16.37 | 0.49 |
| Nt LPPS-AAT4_4_5 | 17.85 | 0.70 |
| Nt LPPS-AAT4_4_6 | 16.85 | 0.56 |
| Mean ± sd | 17.02 ± 0.75 | 0.58 ± 0.11 |
| Nt LPPS-AAT4_8_1 | 10.55 | 0.85 |
| Nt LPPS-AAT4_8_4 | 15.88 | 0.64 |
| Nt LPPS-AAT4_8_6 | 6.49 | 0.46 |
| Mean ± sd | 10.97 ± 4.71 | 0.65 ± 0.20 |

**Table 4.**
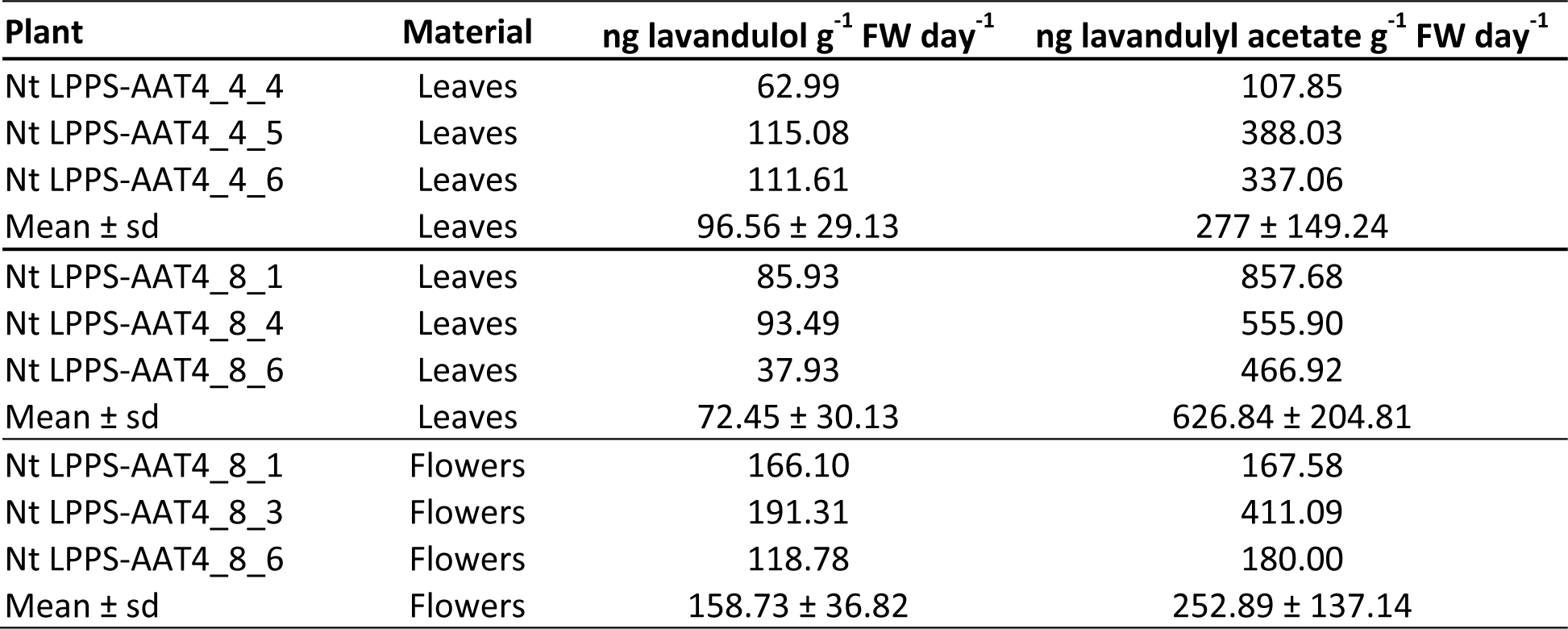
Quantity (ng) of lavandulol and lavandulyl acetate released from organs (leaves and flowers) of T_1_ *NtLPPS-AAT4* plants, obtained by volatile collection and GC/MS/MS quantification.

| Plant | Material | ng lavandulol g<sup>-1</sup> FW day<sup>-1</sup> | ng lavandulyl acetate g<sup>-1</sup> FW day<sup>-1</sup> |
| --- | --- | --- | --- |
| Nt LPPS-AAT4_4_4 | Leaves | 62.99 | 107.85 |
| Nt LPPS-AAT4_4_5 | Leaves | 115.08 | 388.03 |
| Nt LPPS-AAT4_4_6 | Leaves | 111.61 | 337.06 |
| Mean ± sd | Leaves | 96.56 ± 29.13 | 277 ± 149.24 |
| Nt LPPS-AAT4_8_1 | Leaves | 85.93 | 857.68 |
| Nt LPPS-AAT4_8_4 | Leaves | 93.49 | 555.90 |
| Nt LPPS-AAT4_8_6 | Leaves | 37.93 | 466.92 |
| Mean ± sd | Leaves | 72.45 ± 30.13 | 626.84 ± 204.81 |
| Nt LPPS-AAT4_8_1 | Flowers | 166.10 | 167.58 |
| Nt LPPS-AAT4_8_3 | Flowers | 191.31 | 411.09 |
| Nt LPPS-AAT4_8_6 | Flowers | 118.78 | 180.00 |
| Mean ± sd | Flowers | 158.73 ± 36.82 | 252.89 ± 137.14 |

## Discussion

In this study, we demonstrated the potential of tobacco and *N. benthamiana* plants as biofactories of irregular monoterpenes and their derivatives. Several reports show the heterologous production of mono- and sesquiterpenes in bacteria, yeasts and plants (Q. Wang *et al*., 2019; Xie *et al*., 2019; Dusséaux *et al*., 2020; Fuentes *et al*., 2016). However, very few examples exist of the production of irregular monoterpenoids, and many of their biosynthetic pathways remain unknown (Minteguiaga *et al*., 2023). The most widely studied among irregular monoterpene derivatives is chrysanthemic acid, for its importance as the monoterpene moiety of type I pyrethrins. Its full biosynthetic pathway was elucidated (Xu, Moghe, *et al*., 2018) and heterologous expression was achieved in fruits and glandular trichomes of tomato (Xu, Lybrand, *et al*., 2018; Wang *et al*., 2022). Yang *et al*. (2014) also expressed *Tc*CPPS in tobacco to demonstrate its ability to produce chrysanthemol *in planta*, but this was done mainly as a proof of concept rather than to estimate the yield of tobacco biofactories. Concomitantly, heterologous production of insect pheromones has focused almost exclusively on moth sex pheromones, with a few notable exceptions, like the production of 8-hydroxygeraniol in engineered yeast (H. Wang *et al*., 2022), and that of the sesquiterpenoid aphid alarm pheromone (*E*)-β-farnesene in wheat, the first crop engineered to release an insect pheromone (Bruce *et al*., 2015). In this respect, our work moves towards filling this niche and establishing plant-based biofactories for irregular monoterpenoids with prospective uses for extraction and formulation of a variety of products, as well as for live emission in greenhouses or the open field. In this line, we focused on understanding the phenotypic effects associated with bioproduction in unspecialized cells following a constitutive expression strategy, as well as in estimating the biosynthetic potential.

Bioproduction of monoterpenes in *Nicotiana* using a constitutive overexpression strategy is not physiologically innocuous. For instance, Yin *et al*. (2017) found early flowering and increased branching when overexpressing the peppermint geranyl diphosphate synthase small subunit in tobacco. Other deleterious effects were observed for different terpenoids, including chlorosis, dwarfism, and a reduction in fertility (Huchelmann *et al*., 2017). Because of the similarity of the phenotypes observed in different reports, cytotoxicity of the new metabolites is considered not to be the only cause, and perhaps plant depletion of its essential terpenoid precursors (IPP/DMAPP) is also playing a role. Many of the observed deleterious effects in our transgenic plants were dose dependent. We observed a reduction in size and leaf biomass of transgenic plants producing lavandulol and lavandulyl acetate, correlating with the greatest reductions in chlorophyll index. In *N. benthamiana*, the reduction in leaf biomass for *NbLPPS* was due mostly to the observed reduction in number of lateral shoots (data not shown). Despite the observed effects, in general plants showing moderate production levels (e.g., chrysanthemol producers in this study) also showed moderate phenotypic effects, suggesting that a balance between volatile productivity and biomass production can be reached. Whether such balance is favorable in technoeconomic terms will depend on the absolute production levels and the concentrations required for achieving a biological effect.

The analysis of productivity across tissues and developmental stages was useful to understand the dynamics of biosynthesis, as well as the design of the best conditions for biofactory use. Metabolite accumulation depends, at least in part, on precursor availability at a specific growth stage and on the activation of competing metabolic pathways (Drapal *et al*., 2021). Developmental information is also crucial to determine at what stage plant tissues should be harvested or used as bio dispenser to maximize product yield. We found that young leaves at the flowering stage are most productive tissue in most instances. We also observed that, at least in the constitutive overexpression strategy followed here, flowers do not provide a special advantage in terms of accumulation or volatile emission. In the light of these results, it could be advisable to grow plants in short cycles, harvesting before flowering to maximize productivity. Also, non-flowering tobacco varieties (Schmidt *et al*., 2020) could be suitable biofactory candidates, especially as bio-emitters, since genetically impeded flowering would not affect bioproduction negatively and would increase the biosafety profile. Other suggested strategies to increase product yields and to reduce pleiotropic effects in heterologous hosts include accumulation in trichomes using specific promoters and transporters (Huchelmann *et al*., 2017). This might not represent an ideal solution for the accumulation of volatile compounds in tobacco, since it does not possess peltate trichomes such as those of aromatic plants, but rather, its capitate trichomes are specialized for the secretion on the leaf surface of non-volatile diterpenes and phytoalexins (Tissier *et al*., 2017). While it might be a worthwhile approach to increase volatilization for bioemitters, it is possible that the greater mesophyll biomass still guarantees greater yields using ubiquitous promoters, given that fitness loss is kept within acceptable levels.

As expected, absolute product yields were found to be metabolite dependent. Lavandulol yields were consistently higher than those of chrysanthemol (38- and 58-fold more lavandulol than chrysanthemol was extracted from tobacco and *N. benthamiana* tissues, respectively). Previous reports of heterologous expression of *Tc*CPPS found that chrysanthemol may be glycosylated (Xu, Lybrand, *et al*., 2018) and this could account for these consistently lower levels, as well as, possibly, for the lower fitness loss observed in *NbCPPS* and *NtCPPS* plants. Based on the biomass data and the quantification of monoterpenoids, we could estimate the average production per plant for each transgenic line as 0.3 mg chrysanthemol in *NtCPPS* plants, around 16 mg lavandulol for *NtLPPS* plants, and around 3.4 mg lavandulol from a *NtLPPS-AAT4* plant. For lavandulyl acetate, only 0.20 mg of compound per plant could be extracted. It is important to note that the lavandulyl acetate figure is probably underestimating the actual levels of metabolite accumulated in leaves due to partial losses with toluene extraction method. As pointed out, relative quantifications using ground tissue suggest higher contents: lavandulyl acetate was twice the lavandulol detected in *NtLPPS-AAT4*_8 plants, and also twice the lavandulol detected in *NtLPPS*_1_3_1 plants. Alternative strategies for the extraction of lavandulyl acetate based on different solvents or on collection and subsequent extraction of the emitted volatiles could be assayed. Based on toluene extraction levels, approximately 6 kg of young tobacco leaves (6 plants) would be necessary to produce enough lavandulyl acetate to treat 1 L of water against mosquito larvae at its LC50 (Govindarajan & Benelli, 2016). Altogether, it seems clear that at the extractable yields obtained using constitutive overexpression are currently too low to provide a competitive advantage to alternative production systems.

In contrast with the modest technoeconomic perspectives of the constitutive expression strategy in terms of extraction yields, our data suggests a high potential of the tobacco platform as volatile live biodispensers. For this, it is important to put the data obtained here in the context of pest control strategies conducted in the field with related pheromones. Comprehensive assessments of mating disruption strategies to control the mealybug *Planococcus ficus* using lavandulyl senecioate were recently reported by Daane *et al*. (2020) and Lucchi *et al*. (2019) using field deployed dispensers containing chemically synthetized racemic compounds. Here it was found that significant results in pest control were obtained with dispensers loaded with a total of 4.15 g/ha over a season. Likely, the conclusions drawn in these studies may apply to other mealybugs and other pheromones comprising irregular monoterpene esters. According to our daily emission estimations of lavandulyl acetate in tobacco plants (correcting for leaf age and number of leaves at each stage in an adult plant), a few hundred (200-500) plants per hectare would be sufficient to ensure similar release levels as those reported by Lucchi *et al*. (2019). Depending on the crop to which they would be coupled, this might represent a feasible density. The factors relevant for the effectiveness of pheromone dispensers include a steady emission rate (which can be more important than absolute pheromone concentration) and constant coverage during the season, to ensure pheromone emission during all periods of peak flight activity. Plant bioemitters, in this respect, represent interesting solutions because their emission depends on renewable metabolic resources, and it is not restricted to the initial load of the dispenser. The life cycle of a tobacco plant, for example, is compatible with that of other crops to which it may be coupled as producer of pheromones. Also, all available dispensers use racemic mixtures, only half of which is the active ingredient, while biosynthesis ensures stereospecificity.

In conclusion, we show here that tobacco plants producing irregular monoterpenoids, and particularly lavandulyl acetate, are a valuable model to understand the feasibility of using live pheromone emitter plants as tools for mating disruption. Further improvements might be envisioned to increase productivity and volatilization. However, compared to previous works on plant-based production of lepidopteran pheromones (Mateos-Fernández *et al*., 2021), tobacco plants producing monoterpene esters appear to be a more viable tool for plant-based pest control. Finally, in addition to being biofactories and live emitters, these plants represent a versatile metabolic and genetic tool for the combinatorial assessment of a variety of enzymatic activities (e.g., acyltransferases from different sources) acting upon the constitutively expressed monoterpenoid precursors to yield an array of pheromone compounds and other bioactive molecules.

## Supplementary Data

Supplementary Table S1. List of the GB constructs used or generated in this study.

Supplementary Figure S1. Stable production of lavandulol and chrysanthemol in transgenic *N. benthamiana* and tobacco T0 – T2 plants.

Supplementary Figure S2. Levels of Artemisia alcohol, Yomogi alcohol and santolinatriene detected in transgenic *TcCPPS N. benthamiana* and tobacco T0 – T2 plants.

Supplementary Figure S3. Chlorophyll Index (C.I.) of transgenic *N. benthamiana* and tobacco plants producing irregular monoterpenoids.

Supplementary Figure S4. Production of volatile monoterpenoids in transgenic *N. benthamiana* T3 lines in flowers at different development stages.

Supplementary Data File S1. All data and statistical analysis relative to figures in the main text. Supplementary Data File S2. All data and statistical analysis relative to supplementary figures.

## Acknowledgments

We thank Prof. Heribert Warzecha and Elisabeth Haumann from the Technical University of Darmstadt (TUDA), for giving us the sequences of *LiLPPS* and *TcCPPS*. We also wish to thank Dr. Ana Espinosa-Ruiz and Teresa Caballero-Vizcaino at the IBMCP metabolomic facility for their help with GC-MS assays.

## Author contributions

RMF: conceptualization, investigation, writing – original draft preparation; SV: investigation, writing – review and editing; INF: investigation, writing – review and editing; VNL: investigation, writing – review and editing; DO: conceptualization, funding acquisition, writing – original draft preparation; SG: conceptualization, investigation, supervision, writing – original draft preparation. All authors read and approved the final text.

## Data availability

All data reported in this study, as well as statistical analyses, is available in Supplementary Files S1 and S2, deposited at Zenodo: https://doi.org/10.5281/zenodo.8208703. The sequences of the plasmids used for transformation can be consulted at www.gbcloning.upv.es.

