## Supplementary Figure S1-S5 and Supplementary Table S1 for "Assessment of tobacco and *N. benthamiana* as biofactories of irregular monoterpenes for sustainable crop protection"

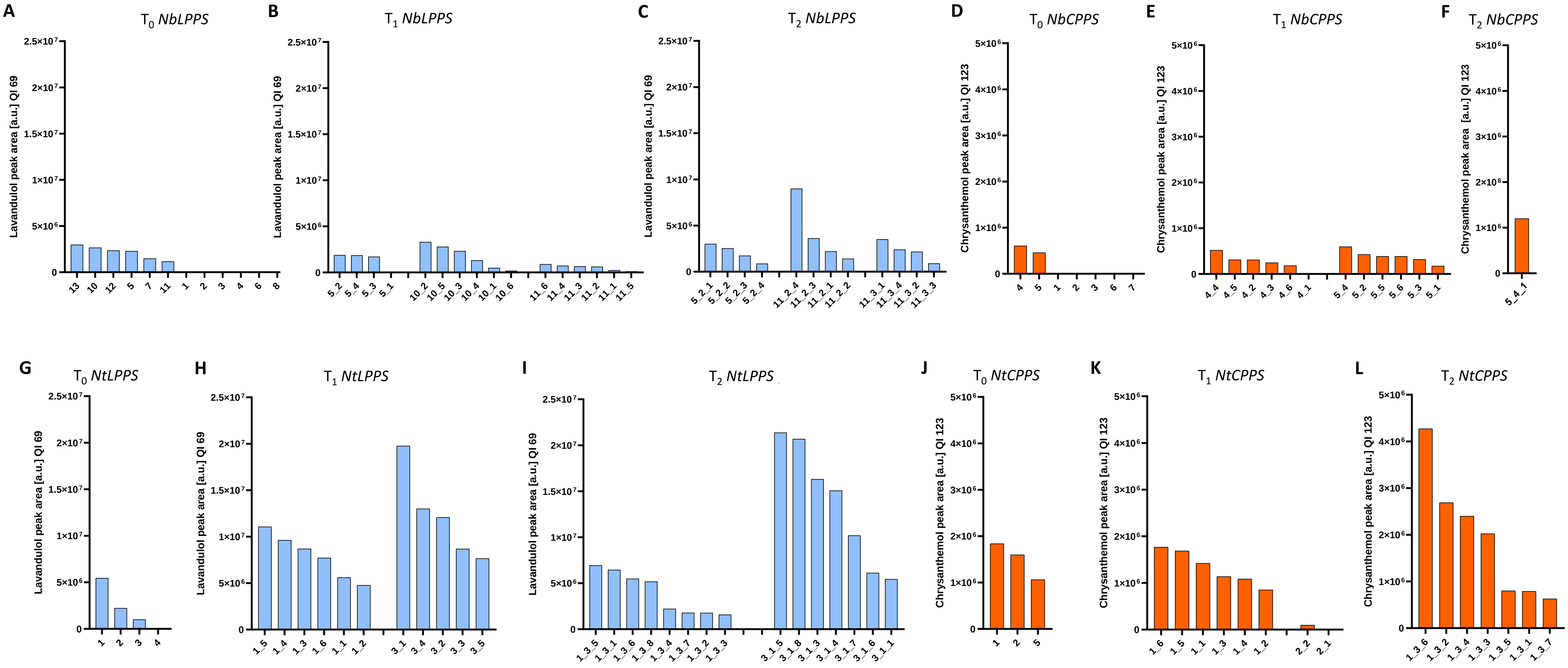


**Supplementary Figure S1. Stable production of lavandulol and chrysanthemol in transgenic *N. benthamiana* and *N. tabacum* $T_0$ – $T_2$ plants.** Lavandulol production levels in individual *NbLPPS* $T_0$ plants (**A**), $T_1$ plants (**B**), and $T_2$ plants (**C**). Chrysanthemol production levels in individual *NbCPPS* $T_0$ plants (**D**), $T_1$ plants (**E**), and $T_2$ plants (**F**). Lavandulol production levels in individual *NtLPPS* $T_0$ plants (**G**), $T_1$ plants (**H**) and $T_2$ plants (**I**). Chrysanthemol production levels in individual *NtCPPS* $T_0$ plants (**J**), $T_1$ plants (**K**), and $T_2$ plants (**L**). Each value represents the production of the mix of the three youngest fully expanded leaves of 35-40 days old plants.

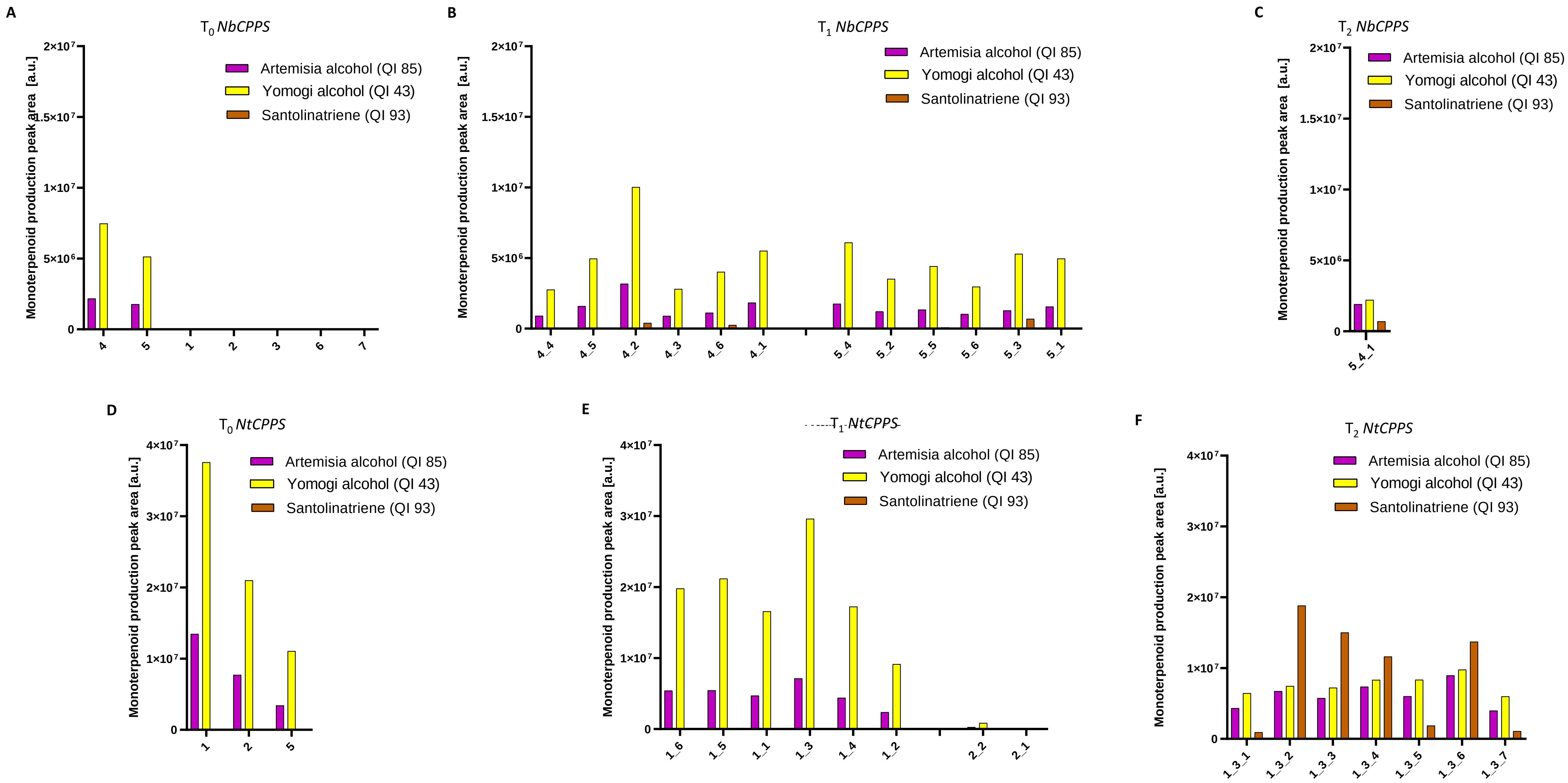


**Supplementary Figure S2. Stable production of artemisia alcohol, Yomogi alcohol and santolinatriene in transgenic *Tc*CPPS *N. benthamiana* and *N. tabacum* $T_0 - T_2$ plants.** Artemisia alcohol, Yomogi alcohol and santolinatriene levels in individual *NbCPPS* $T_0$ plants (**A**), $T_1$ plants (**B**), and $T_2$ plants (**C**). Artemisia alcohol, Yomogi alcohol and santolinatriene production levels in individual *NtCPPS* $T_0$ plants (**D**), $T_1$ plants (**E**) and $T_2$ plants (**F**). Quantification Ions (QI) selected for peak area integration are specific for each molecule, making areas are not fully comparable between compounds. Each value represents the production of the mix of the three youngest fully expanded leaves of 35-40 days old plants.

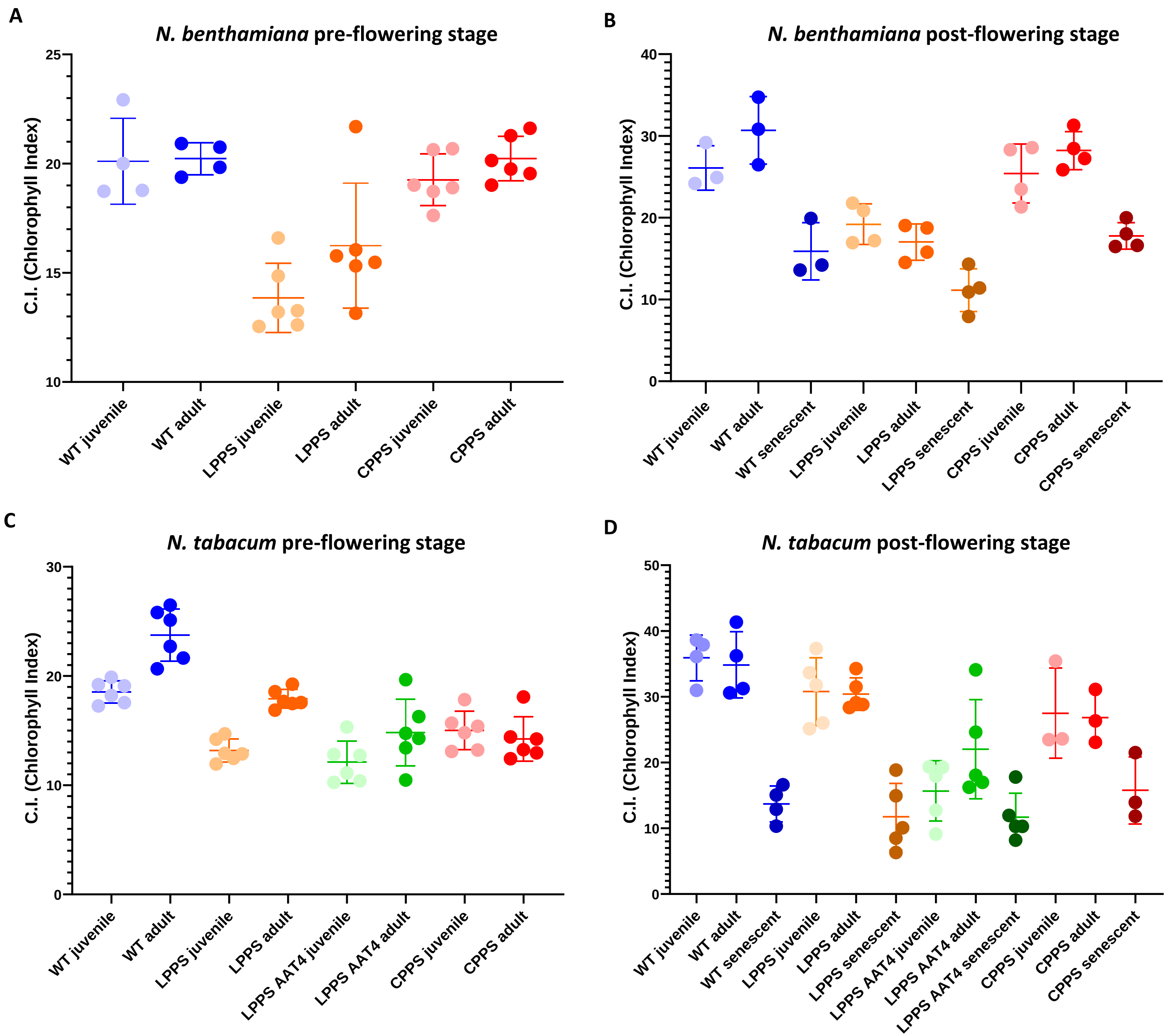


**Supplementary Figure S3. Chlorophyll Index (C.I.) in transgenic *N. benthamiana* and *N. tabacum* plants producing irregular monoterpenoids. (A, B)** Chlorophyll Index measured in juvenile, adult and senescent leaves of $T_3$ *NbLPPS* and *NbCPPS,* and WT *N. benthamiana* in the pre-flowering (A) and post-flowering (B) stages. **(C, D)** Chlorophyll Index measured in juvenile, adult and senescent leaves of $T_3$ *NtLPPS* and *NtCPPS,* $T_1$ *NtLPPS-AAT4* and WT *N. tabacum* in the pre-flowering (C) and post-flowering (D) stages. Values represent the mean and standard deviation of n = 3-6 biological replicates.

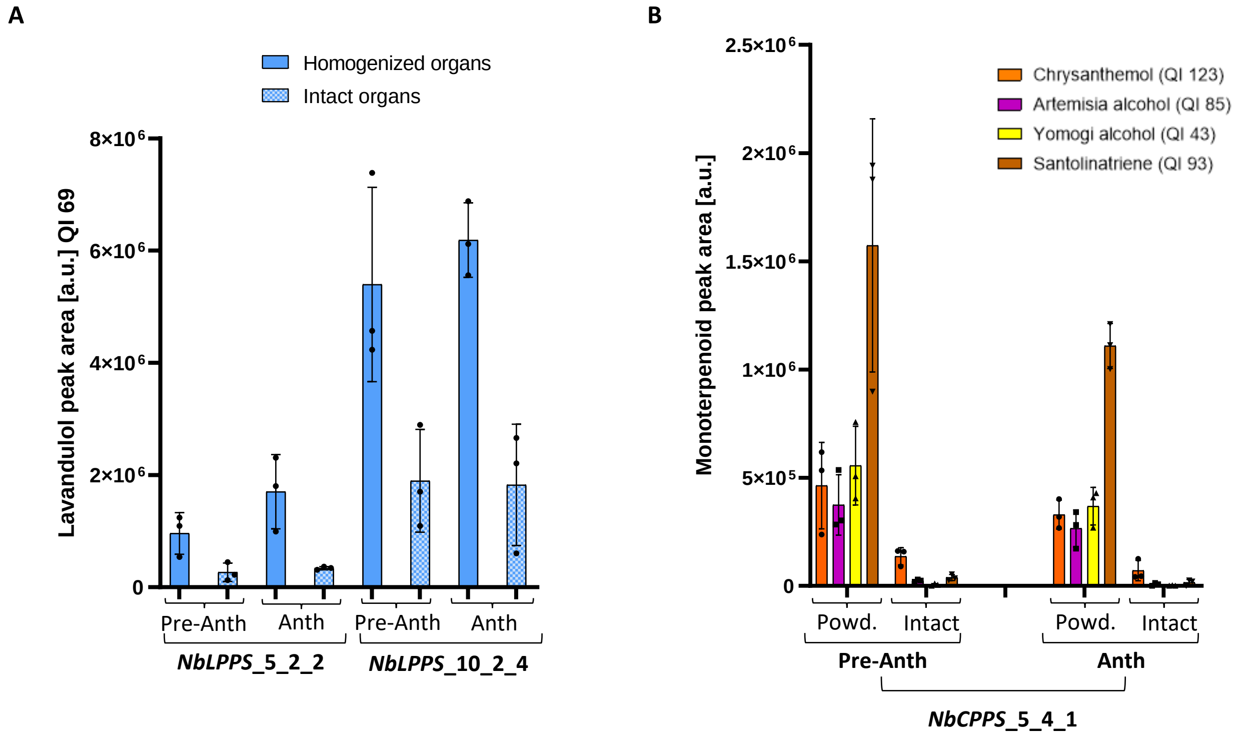


**Supplementary Figure S4. Production of volatile monoterpenoids in transgenic *N. benthamiana* $T_3$ lines in flowers at different development stages**. **(A)** Lavandulol detected in ground samples or emitted from intact flowers of two $T_3$ *NbLPPS* lines, at the pre-anthesis and anthesis stages. **(B)** Chrysanthemol and derived compounds detected in ground samples or emitted from intact flowers of the $T_3$ *NbCPPS*_5_4_1 line, at the pre-anthesis and anthesis stages. Values represent the mean and standard deviation of n = 3 biological replicates (independent flowers). Quantification Ions (QI) selected for peak area integration are specific for each molecule, thus making areas are not directly comparable between compounds.

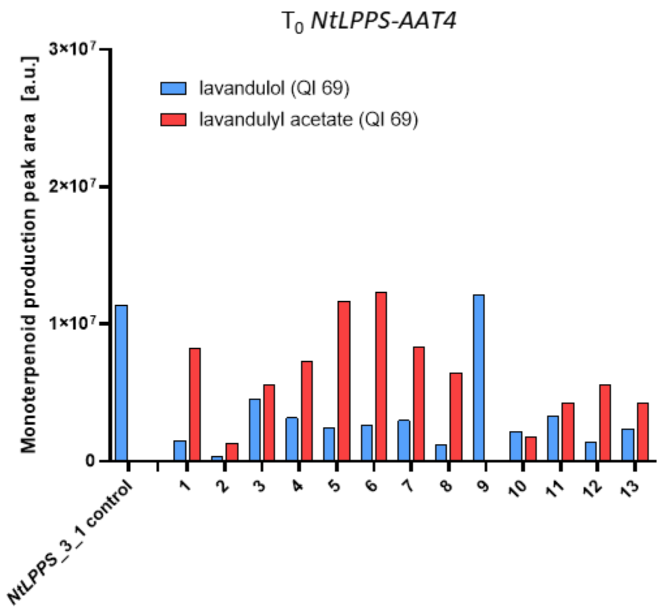


**Supplementary Figure S5. Esterification of lavandulol to lavandulyl acetate in $T_0$ stable tobacco plants**. Production of lavandulol and lavandulyl acetate in $T_0$ *LPPS-AAT4* tobacco plants. Each value represents mixed leaf tissue from the three youngest fully expandend leaves of 35-40 days old plants.

**Table S1**: Existing GoldenBraid constructs used in this study (in blue) and novel GoldenBraid constructs assembled in this study (in black).

| GB ID | Name | Description |
|---|---|---|
| GB0030 | pUPD:p35S | CaMV 35S promoter |
| GB0037 | pUPD:tNos | *Agrobacterium tumefaciens terminator* NOS |
| GB0107 | pEGB SF | Twister plasmid to swap inserts from an alpha1 or alpha1R vector to any omega level vector |
| GB0108 | pEGB p35s:P19:tNos | TU for the expression of the silencing supressor P19 |
| GB0235 | pEGB 1Alpha1R tNos:HygroR:pNos | Hygromycin TU for stable transformation |
| GB1181 | pEGB 3Ω1 tNos:nptII:pNos-SF | TU for kanamycin resistance gene (*nptII*) plant expression under the regulation of the Nos promoter |
| GB3059 | pUPD_CPPS (B4) | pUPD containg CDS of chrysantemyl pyrophosphate synthase from *Tanacetum cinerariifolium* |
| GB3060 | pUPD_LPPS (B4) | pUPD containing CDS of lavandulyl pyrophosphate synthase *from Lavandula x intermedia* |
| GB3061 | 3α1 p35s:CPPS-His:tNos | TU for the expression of chrysanthemyl pyrophosphate synthase (CPPS) with 8xHis |
| GB3062 | 3α1 p35s:LPPS-His:tNos | TU for the expression of lavandulyl pyrophosphate synthase (LPPS) with 8xHis |
| GB3418 | 3Ω2 p35s:LPPS-His:tNos-SF | TU for expression of lavandulyl pyrophosphate synthase (LPPS) with a Stuffer Fragment (SF) |
| GB3419 | 3Ω2 p35s:CPPS-His:tNos-SF | TU for expression of chrysanthemyl pyrophosphate synthase (CPPS) with a Stuffer Fragment (SF) |
| GB3420 | 3α1 p35s:LPPS-His:tNos-SF-tNos:nptII:pNos-SF | TU for expression of lavandulyl pyrophosphate synthase (LPPS) with a Stuffer Fragment (SF) with kanamycin resistance gene (*nptII*) |
| GB3421 | 3α1 p35s:CPPS-His:tNos-SF-tNos:nptII:pNos-SF | TU for expression chrysanthemyl pyrophosphate synthase (CPPS) with a Stuffer Fragment (SF) with kanamycin resistance gene (*nptII*) |
| GB4092 | pUPD2 LiAAT4 | CDS of monoterpene acetyltransferase of *Lavandula x intermedia* LiAAT4 |
| GB4093 | 3α2 p35s:LiAAT4:tNos | TU containing LiAAT4 monoterpenoid acetyltransferase from *Lavandula x intermedia* |
| GB4157 | 3Ω1 tNos:HygroR:pNos p35s:LiAAT4:tNos | Module for expression of LiAAT4 monoterpenoid acetyltransferase from *Lavandula x intermedia* with hygromycin resistance gene |
